# Steering the chain-elongating microbiome to specific medium-chain carboxylates with ethanol and lactate as co-electron donors: maximizing C8 or C6

**DOI:** 10.1101/2025.02.07.637006

**Authors:** Han Wang, Byoung Seung Jeon, Andres E. Ortiz-Ardila, Largus T. Angenent

## Abstract

Microbial chain elongation is a sustainable process to convert organic residues into valuable biochemicals *via* anaerobic fermentation. The operating conditions for the bioreactor and the ecological interactions among functional populations are crucial in controlling this process, but they have not been completely ascertained. Here, we unraveled two operating conditions (*i.e.*, environmental factors): **(1)** the substrate ratio of ethanol and lactate as co-electron donors, and **(2)** the temperature, which affected both the product specificity (*i.e.*, function) and microbial dynamics of chain-elongating microbiomes in a continuously fed bioreactor with product extraction. Specifically, we found that the increase in the substrate ratio of ethanol to lactate shifted the microbiomes toward *n*-caprylate (C8) production, while the slightly higher operating temperatures of 37°C or 42°C were advantageous to *n*-caproate (C6) production. We detected a core microbiome that was similar for all environmental conditions and the two bioreactors, consisting of populations from *Sphaerochaeta* spp., *Caproiciproducens* spp., and Oscillospiraceae. Besides the core microbiome, we observed positive correlations between Erysipelaclostridiaceae UCG-004, *Bacteroides* spp., Oscillospiraceae NK4A214, Rikenellaceae RC9, and *Pseudoclavibacter* spp. with *n*-caprylate production. Similar populations compared to the core microbiome were positively correlated with *n*-caproate production. We showed that we can steer microbiomes toward a high specificity of certain medium-chain carboxylates.

**Abstract Graphics:** 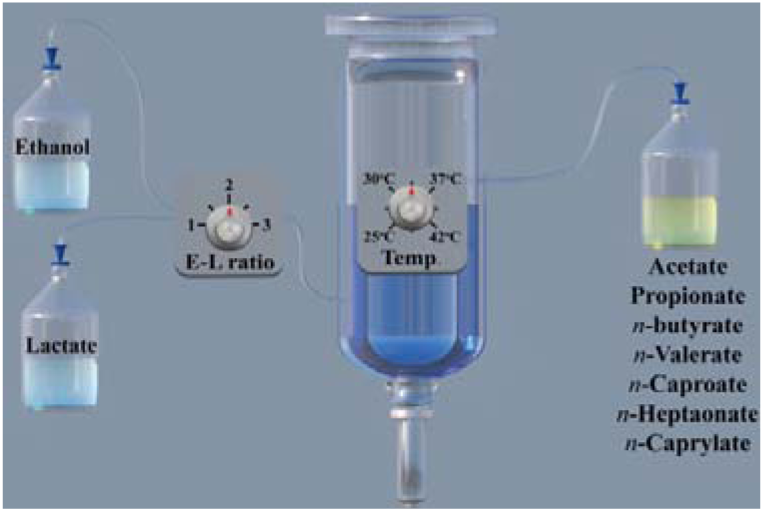

**SYNOPSIS:** Generating medium-chain carboxylates is a promising open-culture biotechnology production platform for converting organic waste streams into biofuels and chemicals.

## 1. INTRODUCTION

The production of synthetic chemicals contributes considerably to current environmental challenges such as global warming, air pollution, and deforestation.^1–3^ Removing the dependency on fossil fuels for chemical commodity synthesis and achieving net-zero CO_2_ emissions would help to alleviate these challenges.^4,5^ When combined with an efficient waste management strategy, a circular economy approach aims to re-value and recover the organic carbon content in waste to produce valuable chemical commodities.^6,7^ Microbial chain elongation to produce medium-chain carboxylates (MCCs), which is based on the open-culture conversion of organic waste into valuable industrial compounds, is considered one promising strategy within the circular economy.^7,8^

MCCs are straight-chain carboxylates with 6-12 carbon atoms (*e.g., n*-caprylate [C8] and *n*-caproate [C6]) and are considered essential chemical commodities due to their multiple applications.^7^ Here, we use the terminology for the dissociated form for all carboxylates to mean the total of the dissociated and undissociated forms. MCCs can be building blocks for liquid biofuels, commercial chemicals, antimicrobials, and feed additives.^9,10^ Microbial chain elongation is performed by the reverse ß-oxidation (rBOX) pathway, which involves two main steps.^11–13^ First, electron donors (*e.g.,* ethanol, lactate, or carbohydrates) are oxidized to generate energy, reducing equivalents, and acetyl-CoA. Second, the electron acceptors (*e.g.,* acetate or propionate) are reduced and elongated with two additional carbon atoms in the form of acetyl-CoA *via* a cyclic pathway.^13^ For example, acetate can be elongated into *n*-butyrate, *n*-caproate, and *n*-caprylate (even-chain carboxylates),^14^ while propionate can be elongated into *n*-valerate and *n*-heptanoate (odd-chain carboxylates).^15^ Ethanol and lactate are the most common electron donors for microbial chain elongation.^16,17^ Promising yields and volumetric production rates of MCCs from ethanol or lactate have been described in chain-elongation bioreactors using pure cultures (*e.g., Clostridium kluyveri*^18^ or Ruminococcaceae bacterium CPB6^19^) and open cultures.^12,14,20,21^

The effluents with organic waste from fermentation-based industries (*e.g.,* acid whey, maize silage, and food waste) may already contain both ethanol and lactate due to the fermentation process and/or the natural presence of lactic acid bacteria during storage.^22,23^ For example, ethanol and lactate were used as co-substrates in producing MCCs from liquor-making wastewater.^24,25^ This combination of electron donors was found to enhance the conversion of SCCs into MCCs.^24,25^ Additionally, the ratio of ethanol to lactate in the substrate was shown to change the product selectivity.^26^ For example, it was reported that the ratio of ethanol to lactate of 2:1 resulted in the highest *n-*caproate selectivity, compared to other ratios of ethanol to lactate.^26^ Furthermore, lactate in the substrate can be converted to propionate, which is the electron acceptor for odd-chain MCC production.^27,28^ The proportion of the odd-chain carboxylates can be controlled by the lactate concentration in the broth.^27^ Also, the ratio of ethanol to lactate in the substrate determined the distribution of even- to odd- chain MCCs that were produced.^26^ However, information on the optimal ratio of ethanol to lactate is scarce during long-term operating conditions, and optimization of this parameter has yet to be systematically investigated.^29,30^

Other operating parameters, such as pH, temperature, and hydraulic retention time, are pertinent for microbial MCC production.^31^ Temperature, for example, is a crucial parameter that influences the thermodynamics and kinetics of metabolic processes.^32,33^ It has been commonly reported that mesophilic temperatures (*e.g.,* between 30°C and 40°C) are more suitable for MCC production^7,8,16^, although this is changing with carbohydrates as electron donors.^34^ Moreover, MCC production in a bioreactor with liquor-making waste increased when the temperature rose from 35°C to 40°C.^35^ A recent study found that the lower range of mesophilic conditions (25°C) was advantageous for *n*-caprylate production at a pH of 7, with activated sludge as a substrate at low production rates.^36^ At the same time, different substrate-based chain elongation processes had various temperature adaptabilities. For example, thermophilic conditions (*e.g.,* 55°C) have been tested without high MCC production rates with ethanol as an electron donor but without *n*-caproate production.^37^ However, polymeric carbohydrates (*e.g.,* starch and hemicellulose) can be converted into *n*-caproate at concentrations up to 283 mg L^-1^ at 55°C.^38^ In addition, the *n*-caproate concentration of 239.7 mmol C L^-1^ (specificity of 40.2%) was obtained from lactate in a complex fermented food waste at 55°C.^39^ Therefore, further studies are required to elucidate the influence of temperature on microbial MCC production using co-electron donors (*e.g.,* ethanol and lactate). Finally, the effects of substrate ratio, pH, and temperature can also shape the reactor microbiome in MCC-producing open cultures.^26,37,40^

Here, we investigated MCC production in open cultures with ethanol and lactate as co-electron donors. The MCC-producing bioreactors were operated in a long-term continuous-run system that was equipped with an in-line extraction module with the specific aim of maximizing *n*-caprylate specificities. We focused on the following three objectives: **(1)** to optimize the use of ethanol and lactate as co-electron donors for MCC production; **(2)** to shape the reactor microbiomes with different ratios of ethanol to lactate and operating temperatures for selective MCC production; **(3)** to investigate the community dynamics by linking the reactor microbiomes with environmental and performance variations. This study provides insight into controlling bioreactor performance and shaping microbiomes for specific MCC production.

## 2. MATERIALS AND METHODS

### 2.1. Inoculum, Bioreactor Setup, and Bioreactor Operating Conditions

Details about the basal medium (**Tables S1-3**), the bioreactor setup (**Fig. S1**), and the in-line extraction system (**Fig. S1**) are given in the Supporting Information. We obtained the inoculum from a long-term operating chain-elongation bioreactor that used ethanol as a substrate.^11,41^ The inoculum was triple-washed in basal media to deplete any remaining substrate and fermentable organic matter before inoculation.^42^ Our operating design included two bioreactors: **(1)** the operating bioreactor (R1); and **(2)** the control bioreactor (R2). For R1, the ethanol-to-lactate ratio (E-L ratio) and operating temperatures were progressively modified, while the R2 was maintained without major modifications, except for Period **c** when the E-L ratio was increased from 1 to 3 during Days 150-190 (**Table S4**). The two bioreactors were supplied with a basal medium that had been previously utilized.^14,27,43^

After inoculation, we fed both bioreactors with ethanol and lactate at a constant total carbon concentration of 695 mmol C L^-1^. We maintained the same conditions during the initial long-term adaptation period (May 2019-June 2020) until both bioreactors performed similarly. Throughout this period, we kept the pH at 5.5, the organic loading rate (OLR) at 110 mmol C L^-1^ d^-1^, the hydraulic retention time (HRT) at 6.4 days, and the E-L ratio at 1 mol mol^-1^. Once both bioreactors reached steady-state conditions, we modified the E-L ratio and operating temperature for R1 (**Table S4**). We varied the E-L ratio between 1 mol mol^-1^ (1:1), 2 mol mol^-1^ (2:1), and 3 mol mol^-1^ (3:1) (**Table S4**), which resulted in concentrations of 139:139 mmol L^-1^, 199:99 mmol L^-1^, and 232:77 mmol L^-1^, respectively, to achieve a constant total carbon concentration of 695 mmol C L^-1^. From here onwards, we will omit the unit mol mol^-1^ for the E-L ratio.

The changes in the operating conditions for R1 were carried out during Periods I-XI. During the first four periods (Periods I-IV), we operated R1 with increasing E-L ratios of 1 (I), 2 (II), and 3 (III), and then with a returned E-L ratio of 1 (IV), while maintaining the temperature at 30°C (**Table S4**). The two Periods V-VI corresponded to a period with a dysfunctional extraction system and a recovery period. Finally, during the five Periods VII-XI, we operated R1 with a temperature gradient of 25°C (VII), 30°C (VIII), 37°C (IX), and 42°C (X), and then with a decreased temperature of 30°C, while maintaining an E-L ratio at 1 (**Table S4**). The operating period for R2 was divided into six periods (**Table S4**): **(1)** Period **a** acted as the control to the operating conditions for R1 during Periods I-IV; **(2)** Period **b** corresponded to a dysfunctional extraction system; **(3)** Period **c** corresponded to the recovery period from Period **b**, including an increase in the E-L ratio (**Table S4**); **(4)** Period **d** acted as the control to the operating conditions for R1 during Periods VII-XI; **(5)** Period **e** corresponded to a failure of the pH probe, resulting in a pH that was lower than 5.5 (approximately 5.0); and **(6)** Period **f** corresponded to the recovery period from Period **e**. Each operating period consisted of at least three HRTs (a 19-day) operating period for R1 and R2.

### 2.2. Liquid Sampling, Analytical Procedures, and Calculations

We collected 1.5-mL samples of bioreactor-mixed liquor every other day from a sampling tube in the middle of the bioreactor height. At the same time, we collected samples from the MCC extraction solution reservoir. All carboxylates were analyzed by gas chromatography (GC) (7890B GC System, Agilent Technologies Inc., Santa Clara, USA), which was equipped with a thermal conductivity detector (TCD) and a capillary column (Nukol Capillary Column: 15 m X 0.25 mm I.D. X 0.25 μm, Agilent Technologies Inc). Ethanol and lactate concentrations were measured using a high-performance liquid chromatography (HPLC) system (Shimadzu LC 20AD, Kyoto, Japan), which was coupled with a refractive index and UV detector (Shimadzu). We also collected and characterized biogas samples daily from the headspace, using GC (SRI gas GCs, SRI Instruments, California, USA). H_2_ and CO_2_ contents were determined by thermal conductivity detector-gas chromatography (GC-TCD), and CH_4_ content was determined with flame ionization detector-gas chromatography (GC-FID). Information on analysis and calculations is given in the Supporting Information.

### 2.3. Biomass Sampling, Sequencing, and Microbiome Analysis

The microbial analysis was based on high-throughput sequencing of bacterial 16S rRNA gene amplicons. Biomass samples were taken from the mixed liquor of the two bioreactors at 56-time points throughout the operating period of ∼1.3 years (in total 112 samples). Each sample was collected in a 2-mL Eppendorf tube with a centrifuge (Eppendorf AG, 22331 Harburg, Germany) at 4°C and 16,873 g for 4 min, while the supernatant was discarded. The pelleted biomass samples were stored at −80 ± 1°C until further processing. Genomic DNA was extracted using the FastDNA™ SPIN Kit for Soil (MO BIO Laboratories Inc., Carlsbad, CA), according to the manufacturer’s instructions. The V4/V5 regions (515F to 926R) of the 16S rRNA gene were amplified from the extracted genomic DNA, following the previously described protocol with slight modifications.^14,44,45^ Amplicon library preparation was performed using dual indexing with a Nextera XT Index kit V2 (Illumina inc., San Diego, USA). Sequencing with the Illumina MiSeq platform was performed at the Max Planck Institute of Biology, Tübingen (Germany) with the MiSeq Reagent Kit v2 (500 cycles).

The obtained sequences were processed with QIIME2.^46,47^ Demultiplexing was performed using the QIIME2 default pipeline, and quality filtering sequence joining, chimera removal, and general denoising were performed using the Divisive Amplicon Denoising Algorithm (DADA2).^48^ After adapter trimming and joining paired reads, 12,058,446 sequences were obtained for the selected samples. This process resulted in 3,001 operational taxonomic units (OTUs) with at least two reads. Taxonomic classification was performed using the machine learning Scikit-learn naive-Bayes classifier^49,50^ with the Silva 138_99 database,^51,52^ setting an 85% acceptance as a cut-off match identity with the obtained OTUs.

α-Diversity metrics, such as the Shannon diversity index and Chao1, were analyzed using QIIME2 and R (version 4.1.3).^47,53^ The Bray-Curtis^54,55^ and unweighted and weighted UniFrac distance metrics^56^ were also obtained from QIIME2. Unless otherwise specified, all plots were generated with ggplot2 package in R.^57^ For this work, community dissimilarities were compared with permutational multivariate analysis of variance (PERMANOVA) *via* the vegan package.^58^ ANOVA, Kruskal–Wallis test, and pairwise Wilcoxon rank sum test were used to identify significant differences between treatments. The normality assumption of the data was tested using the Shapiro–Wilk test on the residuals. P-values were adjusted using the Benjamini–Hochberg correction method. All the statistical analyses were carried out in R.

The β-diversity was determined using distance matrixes obtained from phylogenetic reconstructions carried out in QIME2 and assessed a distance-based redundancy analysis (dbRDA). Non-metric multidimensional scaling (NMDS) analysis^59^ and principal coordinate (PCoA) analysis^60^ were used to visualize the differences in the community among the samples with bray-Curtis or Unifrac distance *via* the vegan package. Only significant (PERMANOVA p<0.001) environmental variables were used as constraints (*i.e.,* ethanol-to-lactate, temperature), correlating the change in the microbial ecology with the MCCs product (*i.e.,* C6 as *n*-caproate and C8 as *n*-caprylate) as eigenvectors. K-means cluster analysis was performed using the phylogenetic distances between populations in the Bray-Curtis matrix present in the microbiome to identify clusters with similar ecological composition (ANOVA p<0.05) within the community by the Stats package for R. Then, using a direct gradient analysis (DGA), significant microbes were ordinated with the constraints of the dbRDA analysis. dbRDA, DGA, clustering calculations, and plots were performed with the Vegan community ecology package.^58^

We utilized the Spearman’s rank correlation coefficient to ascertain whether the microbiomes were significantly correlated or not to MCC production *via* the psych and stats packages.^61^ Heat maps were created using the Ampvis2 package to visualize the abundance of OTUs in different sample points and operating stages.^62^ Finally, we calculated the Spearman’s rank correlation coefficient based on the relative abundances of each OTU. Only OTUs with > 1% relative abundance in more than three samples were included in the analysis. Only the populations with Spearman’s rank correlation coefficient 0.75≤ρ≤1 and FDR-corrected p-value≤0.05 were used for the creation of the eigenvectors.

### 3. RESULTS AND DISCUSSION

### 3.1. Bioreactor Performance

#### 3.1.1. A Higher E-L Ratio Benefited *n*-Caprylate Production and Even-chain Carboxylate Production

We varied the E-L ratio between 1 and 3 for R1 to determine the effect on MCC production (Periods I-IV, **Fig. 1A-C**). The E-L ratio for R2 during Period **a** was maintained at 1 as the control (**Fig. 1D**), which led to a stable product spectrum, with *n*-caproate at an average volumetric production rate of 42.5 mmol C L^-1^ d^-1^ as the predominant product with an average selectivity (*i.e.*, product divided by substrate) of 0.39 mmol C mmol C^-1^ and average specificity (*i.e.*, product divided by all carboxylate products) of 0.56 mmol C mmol C^-1^ (**Fig. 1D**, **Fig. S2B**, and **Table S5-S6**). Indeed, this result was similar to Period I for R1 with an identical E-L ratio of 1 (47.0 mmol C L^-1^ d^-1^, 0.43 mmol C mmol C^-1^, and 0.54 mmol C mmol C^-1^, respectively, in **Fig. 1A**, **Fig. S2A**, and **Table S5-S6**). The biogas composition for both bioreactors showed a constantly low fraction of H_2_ of 1-3%, which is high enough to prevent syntrophic carboxylate oxidation under anaerobic conditions and low enough to not slow down chain elongation (**Fig. S3-S4**).

**Fig. 1.**
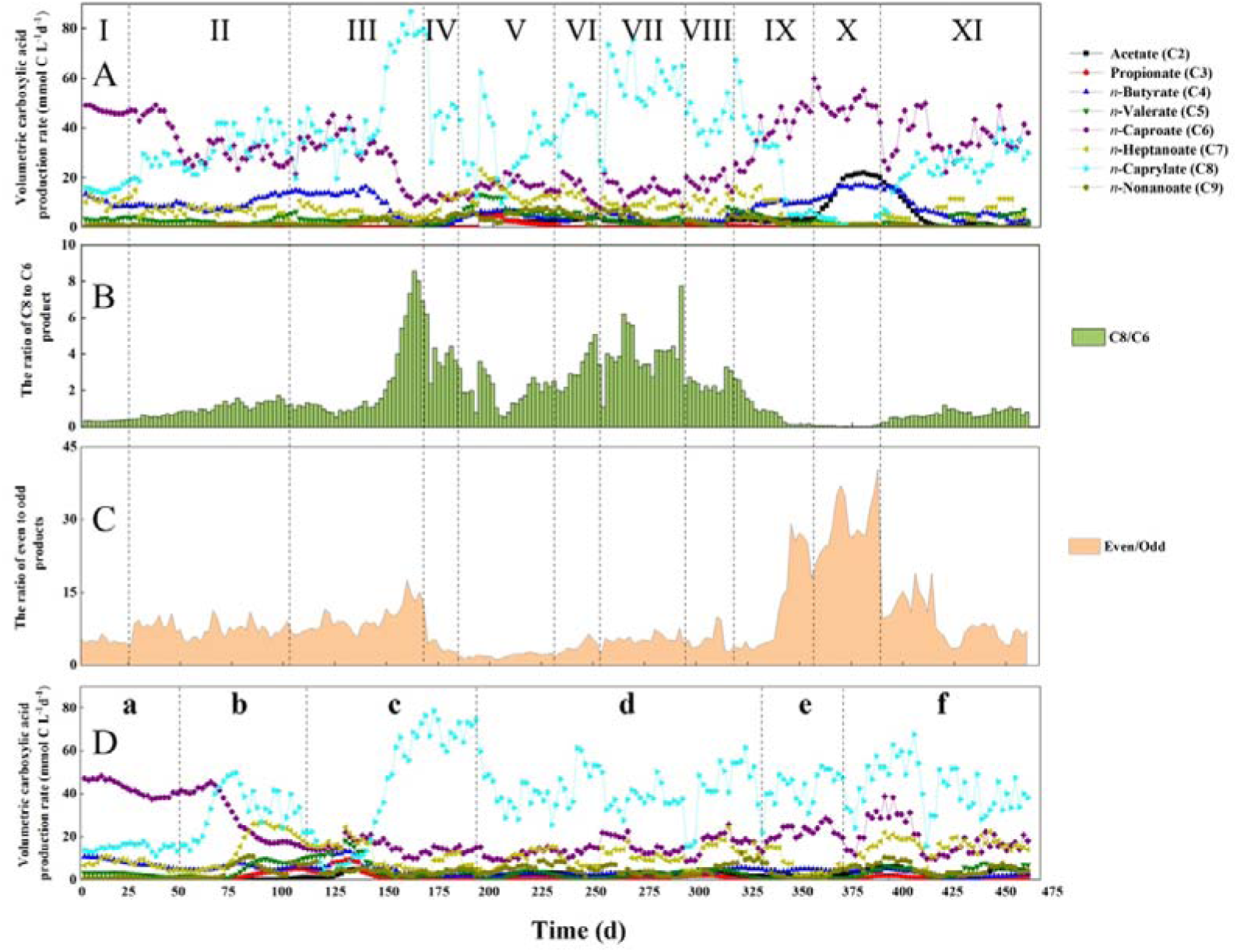
Performance of the bioreactors: (A) line-area chart for the production rates of carboxylates in R1; (B) column chart for the C8/C6 ratio (*i.e.*, *n*-caprylate/*n*-caproate ratio) in R1; (C) stacked chart for the even/odd (*i.e.*, even-/odd-chain carboxylates) in R1; (D) line-area chart for the production rates of carboxylates in R2. The ratios for R2 are shown in **Fig. S5**. The data represent a 6-day moving average.

By increasing the E-L ratio from 1 to 3 for R1, we increased the *n*-caprylate production and the even-chain carboxylate production (Periods I-III, **Fig. 1A-C** and **Table S5**). When we increased the E-L ratio to 2 during Period II, the average volumetric *n*-caprylate production rate increased by 133% during steady-state conditions (from 16.0 to 37.2 mmol C L^-1^ d^-1^ in **Table S5**), while the average volumetric *n*-caproate production rate decreased by 37% (from 47.0 to 29.4 mmol C L_-1_ d^-1^ in **Table S5**). This increased the average *n*-caprylate selectivity and specificity from 0.15 to 0.34 mmol C mmol C^-1^ and from 0.18 to 0.43 mmol C mmol C^-1^ between Period I and II, respectively (**Table S6**). We increased the E-L ratio to 3 during Period III, resulting in *n*-caprylate becoming the dominant product in the bioreactor with an average *n*-caprylate volumetric production rate of 79.3 mmol C L^-1^ d^-1^ (0.06 g L^-1^ h^-1^), and an average *n*-caprylate selectivity and specificity of 0.72 mmol C mmol C^-1^ and 0.78 mmol C mmol C^-1^, respectively, during steady-state conditions, which were the maximum values in this study (**Fig. 1A**, **Fig. S2A**, and **Table S5-S6**). Because *n*-caprylate was extracted at considerably faster rates than *n*-caproate, the average total MCC selectivity and specificity were also the highest in this study during steady-state conditions, with 0.89 mmol C mmol C^-1^ and 0.97 mmol C mmol C^-1^ (**Table S6**). For R2, we increased the E-L ratio from 1 to 3 to recover the control bioreactor and this showed also an increase in the average *n*-caprylate volumetric production rate to 55.7 mmol C L^-1^ d^-1^, and an average *n*-caprylate selectivity and specificity of 0.51 mmol C mmol C^-1^ and 0.89 mmol C mmol C^-1^, respectively, during steady-state conditions (**Fig. 1D**, **Fig. S2B**, and **Table S5-S6**). Our work agrees with Lambrecht *et al.*^63^, who found that ethanol was linked to *n*-caprylate and lactate was linked to *n*-caproate production.

During Periods I-III, the increase in E-L ratio from 1 to 3 was accompanied by an increase in the average ratio of *n*-caprylate to *n*-caproate production during steady-state conditions from 0.34 to 7.56 mol C mol C^-1^, with a maximum exceeding 8 mol C mol C^-1^ on Day 170 (C8/C6 in **Fig. 1B** and **Table S5**). For R2 at an E-L ratio of 3, this maximum ratio reached 6.5 (**Fig. S5A**). According to previous studies, more ATP is released when longer MCCs are produced, which is advantageous for chain-elongating microbiomes.^14^ To date, the production of *n*-caprate (C10) *via* chain elongation has not been reported, which we also did not observe here. Instead, the increased E-L ratio resulted in *n*-caprylate as the dominant MCC at an E-L ratio of 3. We returned the E-L ratio to 1 during Period IV, and this resulted in a 46% decrease in the average *n*-caprylate volumetric production rate to 42.8 mmol C L^-1^ d^-1^ (**Table S5**), while the *n*-caproate volumetric production rate remained fairly constant (**Fig. 1A**), reducing the average ratio of *n*-caprylate to *n*-caproate production from 7.56 to 3.61 mol C mol C^-1^ (**Fig. 1B**). However, we observed hysteresis because this ratio did not return to 0.34 and 0.38 mol C mol C^-1^, which we observed at an E-L ratio of 1 during Period I for R1 and during Period **a** for R2, respectively (**Fig. 1B**, **Fig. S5A**, and **Table S5**).

Throughout Periods I-III, the average ratio of the total concentration of even-chain carboxylates (*i.e.*, acetate, *n*-butyrate, *n*-caproate, and *n*-caprylate) to odd-chain carboxylates (*i.e.*, propionate, *n*-valerate, *n*-heptanoate, and *n*-nonanoate) increased from 4.90 to 13.3 mol C mol C^-1^ (**Fig. 1C**). Both even-and odd-chain MCCs were present in our bioreactor with ethanol and lactate as co-electron donors. First, acetate, which is an electron acceptor for even-chain carboxylate production, can be produced from ethanol and lactate.^64^ Second, propionate, which is an electron acceptor for odd-chain MCCs,^15^ can be produced from lactate *via* the Wood-Werkman cycle or acrylate pathway.^27,40^ When we applied an increasing E-L ratio during Periods I to III, we decreased the lactate concentration in the medium for R1. A reduction in the volumetric lactate loading rate innately decreased the carbon flux toward propionate production, resulting in a lower production of odd-chain carboxylates.^27^ Therefore, a higher E-L ratio led to a higher ratio of even-chain to odd-chain carboxylate production (>30 mol C mol C^-1^ in Period I-III, **Fig. 1C**), which was immediately reversed by reducing the E-L ratio again (Period IV, **Fig. 1C**). Thus, the manipulation of the E-L ratio can steer the MCC product spectrum.

#### 3.1.2. A Higher Operating Temperature Benefited *n*-Caproate Production and Even-Chain Carboxylate Production

We varied the operating temperatures between 25°C and 42°C at a constant E-L ratio of 1 for R1 to test the effect of operating temperature on MCC production (Periods VII-XI, **Fig. 1A-C** and **Table S5**). The operating temperature was maintained at 30°C for R2 as the control, resulting in relatively stable MCC production conditions throughout Period **d** (**Fig. 1D**, **Fig. S5A-B,** and **Table S5**). At relatively low operating temperatures of 25°C-30°C, *n*-caprylate was the dominant product for R1 (Periods VII and VIII, **Fig. 1A**). In addition, the average volumetric *n*-caprylate production rate at 25°C was 30% higher than at 30°C (59.2 mmol C L^-1^ d^-1^ *vs.* 41.4 mmol C L^-1^ d^-1^ in **Table S5**). This resulted in an average *n*-caprylate selectivity and specificity of 0.54 mmol C mmol C^-1^ and 0.65 mmol C mmol C^-1^, respectively (**Table S6**), with a relatively high total MCC selectivity and specificity of 0.77 mmol C mmol C^-1^ and 0.92 mmol C mmol C^-1^, respectively, during steady-state conditions (**Table S6**). Wang *et al.* also observed a higher *n*-caprylate production at 25°C than at 30°C.^36^ When we increased the operating temperature from 30°C to 42°C, the average volumetric *n*-caprylate production rate decreased further from 41.4 to 0.89 mmol C L^-1^ d^-1^ (**Fig. 1A** and **Table S5**). On the contrary, the volumetric *n*-caproate production rate increased from 18.8 to 48.1 mmol C L^-1^ d^-1^ at 30°C and 42°C, respectively (**Fig. 1A-B** and **Table S5**), resulting in an average *n*-caprylate-to-*n*-caproate ratio of 0.04 mol C mol C^-1^ (**Fig. 1B**). A previous study found that when the temperature was increased from 35°C to 40°C, both *n*-caproate and *n*-caprylate production improved, but we did not observe this result.^35^ A return of the operating temperature to 30°C slowly restored the volumetric *n*-caprylate production to an average of 27.5 mmol C L^-1^ d^-1^ during Period XI (**Fig. 1A** and **Table S5**). Meanwhile, the average *n*-caproate production rate remained relatively high at 35.4 mmol C L^-1^ d^-1^ after an initial drop (**Fig. 1A** and **Table S5**), resulting in an average *n*-caprylate-to-*n*-caproate ratio of 0.80 mol C mol C^-1^ (**Fig. 1B** and **Table S5**).

The above results revealed that the lower operating temperatures favored *n*-caprylate production, while the MCC product spectrum shifted toward *n*-caproate production at higher temperatures. The temperature considerably influences the energy released from reactions and affects the kinetic rates of metabolic reactions of the reactor microbiomes.^32^ It is also possible that *n*-caprylate as a product inhibits cellular growth at higher temperatures, even when still in the mesophilic range. The production of longer chain-length carboxylates (*e.g., n*-caprylate) by the reactor microbiomes was almost completely suppressed at 42°C. Another study found that the intermediate substrates (*e.g.,* acetate and *n*-butyrate) for MCC production would accumulate if they were not used for chain elongation.^13^ Indeed, we observed that the average acetate and *n*-butyrate production rates increased in the bioreactor at 42°C during Period X (**Fig. 1A**).

The consequential accumulation of acetate and some *n*-butyrate during Period X (**Fig. S6**) occurred due to the considerably slower extraction rate for SCCs than MCCs,^11^ resulting in an overall lower extraction rate of the total undissociated carboxylic acids, and thus a lower average base (NaOH) consumption rate for extraction during Period X (**Fig. S7**). In addition, we observed the complete circumvention of pH control for our chain-elongation bioreactor, with an average acid consumption rate of 0 g HCl L^-1^ d^-1^ during Period X with an E:L ratio of 1 (**Fig. S7** and **Table S7**). Roghair already predicted the possibility of circumventing pH control because chain elongation with ethanol produces protons, while chain elongation with lactate consumes protons.^65^ Thus, we circumvented pH control by feeding both ethanol and lactate and then by increasing the operating temperature until the composition of the carboxylates was optimum to not extract too many protons with the undissociated carboxylic acids. When we decreased the temperature from 42°C to 30°C again, the volumetric acetate and *n*-butyrate production rates decreased from 18.5 to 0.74 mmol C L^-1^ d^-1^ and from 15.8 to 4.22 mmol C L^-1^ d^-1^, respectively (Period X-XI in **Fig. 1A** and **Table S5**), resuming acid consumption to 0.06 g HCl L^-1^ d^-1^ by the pH-control system for the chain-elongating bioreactor (**Fig. S7**). In principle, a pH auxostat with lactate as a co-electron donor could control the pH without adding HCl when ethanol is the other co-electron donor.

The average ratio of even- to odd-chain carboxylates increased approximately 450% from 5.61 to 30.6 mol C mol C^-1^ after increasing the operating temperature from 25°C to 42°C during Periods VII-X, indicating that higher operating temperatures promoted the production of even-chain carboxylates (Even/Odd in **Fig. 1C** and **Table S5**). In addition, R1 produced a relatively low average concentration of odd-chain carboxylates at 42°C (**Table S5**). The decrease in total odd-chain MCCs may be due to the inhibition of propionate production at higher operating temperatures. From a thermodynamic feasibility standpoint, we found that the free energy changes (ΔG°’ at pH=5.5) for propionate production became less negative when operating temperatures increased (**Eq. 7** in **Table S8**), which implied that the conversion of lactate to propionate became less feasible at a higher temperature. The reduction in propionate production led to an insufficient flux of electron acceptors toward odd-chain MCC production. Similar to the average ratio of *n*-caprylate to *n*-caproate, reducing the operating temperature from 42°C to 30°C led to an immediate decrease in the ratio of even- to odd-chain carboxylate production with an average ratio of 7.23 mol C mol C^-1^ (from 30.6 mol C mol C^-1^) (**Fig. 1C** and **Table S5**). Thus, the overall odd-chain MCCs decreased with rising temperature from 25°C to 42°C during Periods VII-X. Here, we found that the operating temperature provided a control tool to steer the bioreactor to the target product. Finally, we observed an increase in the ratio of the even-chain to odd-chain carboxylates to approximately 15 mol C mol C^-1^ due to an accidental drop in the pH from 5.5 to approximately 5.0 during Period **e** (**Fig. S5B**).

#### 3.1.3. An Efficient in-Line Extraction System was Crucial to Substrate Utilization and Stable MCC Production with Ethanol and Lactate as Co-Electron Donors

Immediately after increasing the E-L ratio from 1 to 2 (at the start of Period II) or from 2 to 3 (at the start of Period III), residual substrate (*e.g.,* ethanol) was detected in R1 (**Fig. S8A**). However, all substrates were completely consumed again after an adaptation period during Periods II or III (**Fig. S8A**). Similarly, when we increased the E-L ratio from 1 to 3 during Period **c** for R2, residual ethanol was detected in the bioreactor, which was consumed again after an adaptation period (**Fig. S8B**). The microbiomes during Periods II, III, or **c** consumed the increased ethanol availability by producing *n*-caprylate rather than *n*-caproate, because we found an increasing volumetric *n*-caprylate production rate during Periods II, III, or **c** (**Fig. 1A** and **Fig. 1D**). However, a positive correlation exists between the carbon chain length of MCCs and its toxicity, resulting in a more severe product inhibition for *n*-caprylate than for *n*-caproate.^66–68^ Fortunately, we operated an in-line extraction system to extract MCCs at a pH of 5.5, avoiding product inhibition. The extraction system maintained a low concentration of undissociated carboxylic acids in the bioreactor, such as 2.42 mM for undissociated *n*-caproic acid, 0.01 mM for undissociated *n-*heptanoic acid, and 0.38 mM for undissociated *n*-caprylic acid, which are the toxic chemical species (calculated based on average total carboxylate concentrations in **Table S9** from individual datapoints in **Fig. S6**). Our extraction system is selective with a faster extraction rate for *n*-caprylate than *n*-caproate.^41,69,70^

Without the in-line extraction system that extracts more efficiently when the MCCs have a longer chain length, ethanol would have accumulated throughout the periods of elevated E-L ratios in our study. Indeed, a dysfunctional extraction system during Period **b** for R2, resulted in the highest residual ethanol concentration of this study, which exceeded 45 mM (**Fig. S8B**), leading to a relatively low MCC production rate at the end of Period **b** (**Fig. 1D**). This residual ethanol concentration at an E-L ratio of 1 was also much higher than the residual lactate concentration of approximately 5 mM (**Fig. S8B**). In addition, previous research showed ethanol accumulation (and not lactate accumulation) during the biological MCC production process when both ethanol and lactate were present in the substrate.^29,30^ With elevated *n*-caproate concentrations when extraction is absent, the pH was one of the selective factors affecting ethanol or lactate utilization for MCC production.^63^ Due to a narrow preferred pH range, ethanol-based chain elongation is more sensitive to product toxicity than lactate-based chain elongation. In contrast, lactate-based chain elongation produces MCCs at pH levels from 5.5 to 7.0. In a previous study at a pH from 4.5 to 5.5, ethanol accumulated, but lactate was still utilized during a period with an undissociated *n*-caproic acid concentration of 8.3 mM.^30^ This concentration was slightly higher than the 7.5 mM, which was shown to be toxic to ethanol-based chain elongation in another study.^41^ Thus, from previous studies with ethanol and lactate as co-substrates for chain elongation,^29,30^ the conversion of primary lactate as an electron donor may lead to the theory that simultaneous chain elongation is not feasible. However, we dispute this theory with long-term adapted reactor microbiomes and product extraction where ethanol and lactate were utilized simultaneously.

### 3.2. The Substrate Ratio and Operating Temperatures Shaped the Reactor Microbiome for Target MCC Production

Our study revealed dynamic shifts in the ecological structure of the reactor microbiomes in response to changes in environmental conditions. Using distance-based redundancy analysis (dbRDA) (**Fig. 2**) and the heatmaps of the observed clusters (**Fig. S9** and **Fig. S10**), we tracked shifts in microbiome composition throughout the operating periods for R1 and R2, focusing on variations in substrate ratio for R1 and R2 and temperature for R1. The ecological succession of each microbiome (each microbiome sample is a symbol in **Fig. 2**) for R1 throughout the entire operating period was organized into four main clusters, reflecting changes in α diversity. As observed in the dbRDA, the four clusters are organized chronologically as described here and with the corresponding heatmaps in the same order (**Fig. S9A-D**). First, the microbiome samples that grouped in a *red* cluster highlighted a significant correlation between *n*-caprylate production (C8 eigenvector) and high ethanol-to-lactate ratios of 2 and 3 (blue and red dots for Periods II and III) (PERMANOVA < 0.001, **Fig. 2A**). Second, the microbiome samples in the *green* cluster corresponded to almost only periods following acute changes at a temperature of 30°C (orange and red dots), such as a shift in the ethanol-to-lactate ratio from 3:1 to 1:1 (red dots for Period IV), extraction system failures (Period V), and the shift from the high-temperature period to the low-temperature period (end of Period XI) (**Fig. 2A**). Third, the microbiomes that formed a *purple* cluster were sampled almost only during periods of optimal mesophilic growth and recovery from operating changes, with temperatures between 25°C and 30°C (orange dots and triangles in **Fig. 2A**). Fourth, for the microbiome samples that grouped in a *blue* cluster, *n*-caproate production (C6 eigenvector) was correlated to increased temperatures or a transitioning period at the end of the operating period (orange squares for Period X and orange dots for the start of Period XI) (**Fig. 2A**). These changes indicate ecological disturbances that prompted the microbiomes to modify their ecological structure and adapt to new environmental conditions.

**Fig. 2.**
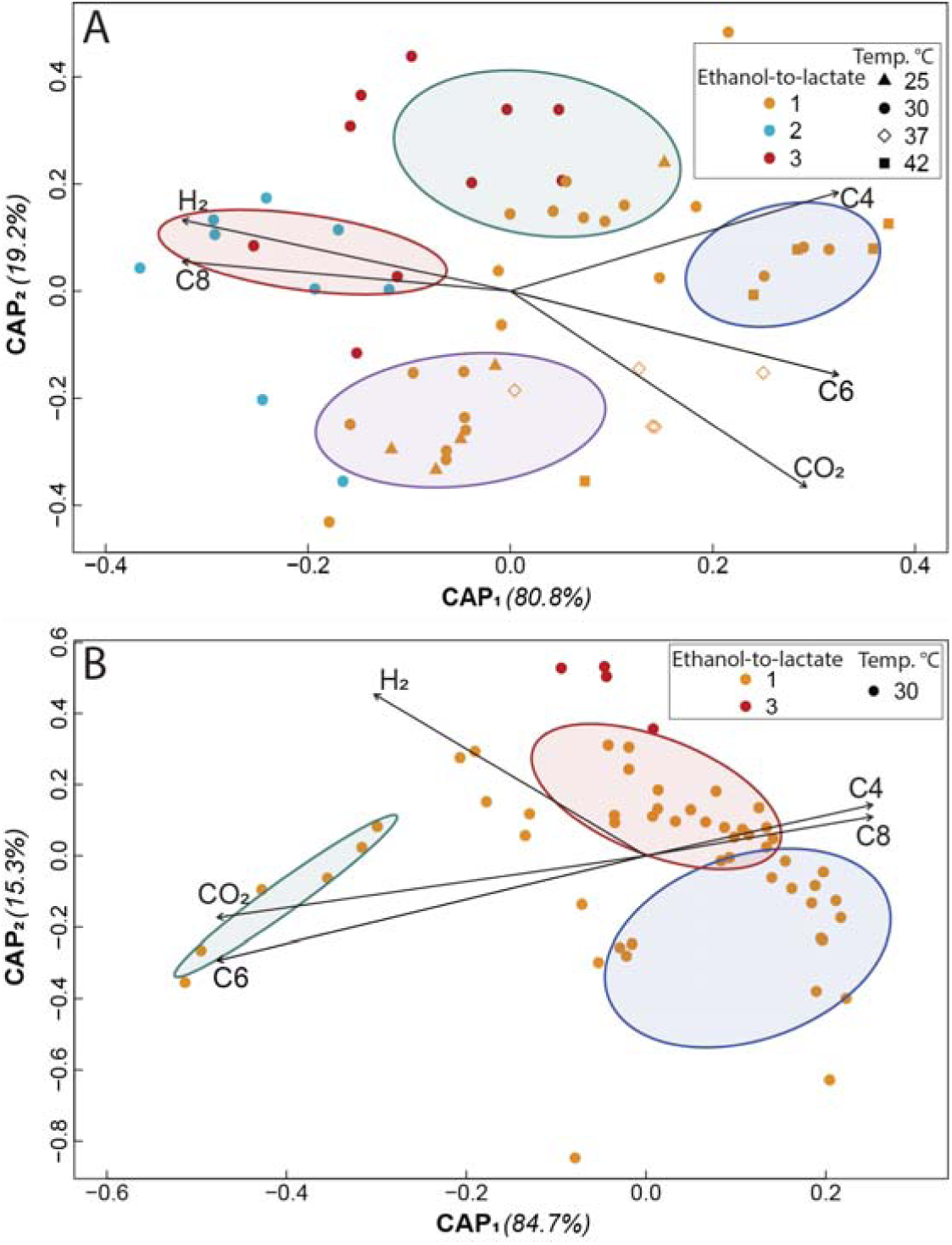
Distance-based redundancy analyses using Bray-Curtis ß-diversity distance matrixes of all microbiome samples during the operating periods for the experimental bioreactor R1 **(A)** and the control bioreactor R2 **(B)**. The eigenvectors were calculated based on PERMANOVA significances < 0.001; and the cluster analysis was performed based on the *k*-means clustering algorithm using square-mean differences between populations in the distance matrixes with significances < 0.05. Cluster colors in A: red (left); green (center up); purple (center down); and blue (right). Cluster colors in B: green (left); blue (right down); and red (right up). Vectors: H_2_ (hydrogen); CO_2_ (carbon dioxide); C4 (*n*-butyrate); C6 (*n*-caproate); and C8 (*n*-caprylate).

Despite these disturbances, for example, in the *green* cluster, three microbial populations from *Sphaerochaeta* spp., *Caproiciproducens* spp., and the Oscillospiraceae family remained consistently dominant across most samples from both bioreactors, forming a core microbiome (**Fig. S9** and **Fig. S10**). The combined relative abundance for these three populations ranged between 42.5% and 75.1% and exhibited resilience, stability, and strong ties to MCC production as the microbiome’s key function for experimental bioreactor R1 (**Fig. S9**). In addition, the resilience of these core populations was similar to control bioreactor R2, where their combined relative abundance remained constant (48.9–67.7%), serving as an indicator of stable environmental conditions (**Fig. S10**). Therefore, these populations met the criteria for a core microbiome, including: **(1)** consistent dominance across samples regardless of operating conditions;^71^ **(2)** resilience and stability; **(3)** close relationship with the main ecological function of the microbiome;^72,73^ and **(4)** serving as indicators for stable environmental conditions.^74^ Indeed, previous studies have shown that these core taxa are closely related to MCC production as the main ecological function of the microbiome.^75–79^ We observed that these populations in the core microbiome were positively correlated with either *n*-caprylate (C8 eigenvector) or *n*-caproate production (C6 eigenvector), which identified that the same core populations produced *n*-caproate for periods (I-IV in **Fig. 3A** and **a-b** in **Fig. 3B**) because *n*-caproate is elongated to *n*-caprylate. Section 3.3 will describe the populations that further elongated *n*-caproate into *n*-caprylate during periods of high *n*-caprylate production (I-IV in **Fig. 3A**). Here, we are not discussing the populations, such as *Pseudoramibacter* spp., that were solely correlated to the C6 eigenvector (**Fig. 3B**).

**Fig. 3.**
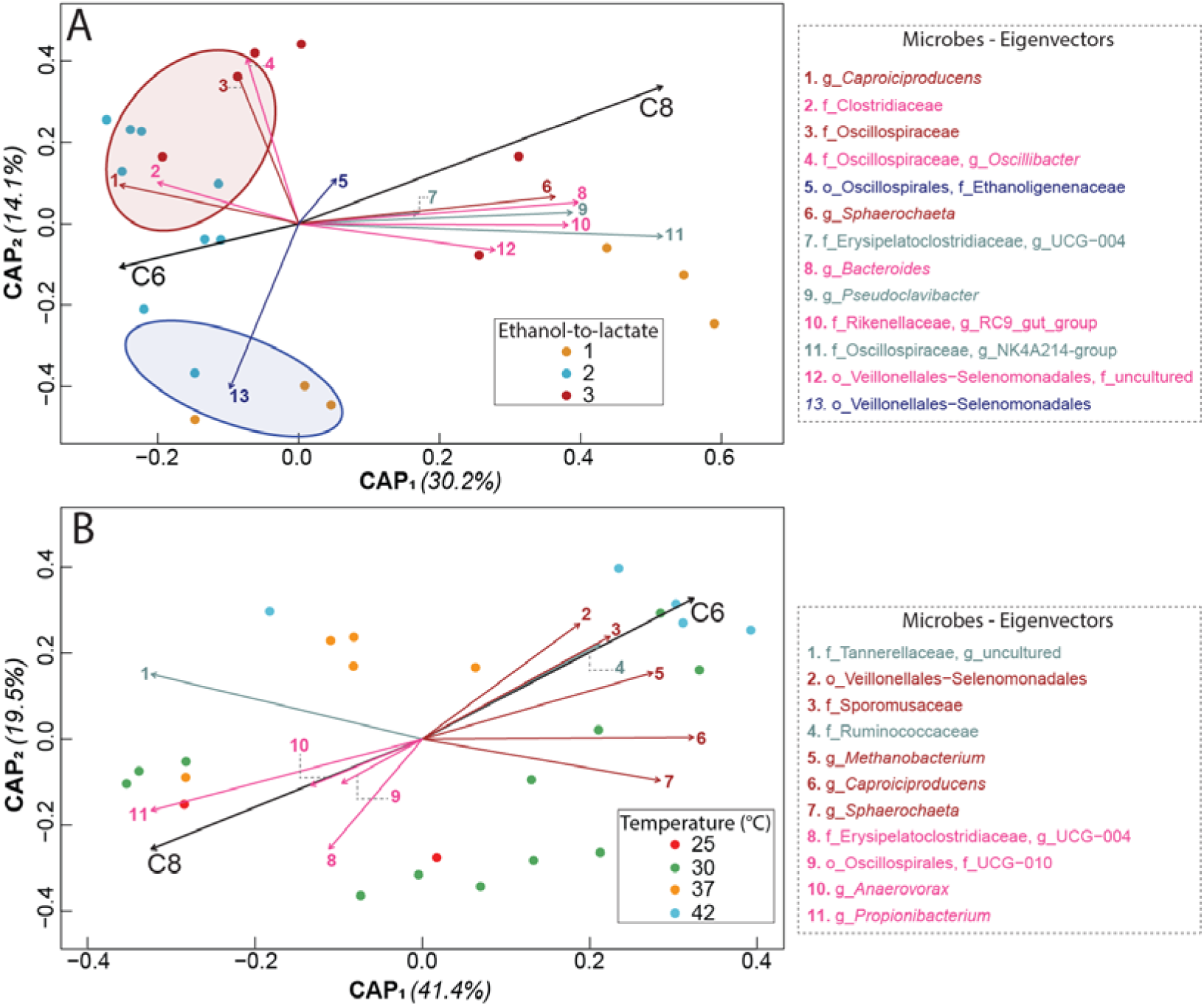
Component analyses for Periods I-IV with different ethanol-to-lactate ratios for R1(**A**) and Periods VII-XI with different temperatures for R1 (**B**) using distance-based redundancy analyses with Bray-Curtis ß-diversity distance matrixes. The eigenvectors were calculated based on PERMANOVA significances <0.05, while the cluster analysis was performed based on *k*-means clustering algorithm using square-mean differences between populations in the distance matrixes with significances <0.05. Eigenvectors: core microbiome members (red); microbes with low significance (blue); significant microbes with correlation coefficients <0.08 (pink); highly significant microbes with correlation coefficients >0.08 (green). Other vectors: C6 (*n*-caproate) and C8 (*n*-caprylate).

For R1, the disturbance in the core population’s abundance correlated with the magnitude of the shifts in operating conditions. For example, we changed the temperature (*i.e.,* periods VII to XI), which caused disturbances in the microbiome composition. When we lowered the temperature from 30°C to 25°C during Period VII in the *purple* cluster, *Caproiciproducens* spp. and Oscillospiraceae populations, which are core members, increased compared to Period VI (**Fig. S9**), while the volumetric *n*-caprylate production rates increased (**Fig. 1A**). Relatively high temperatures also strongly disturbed the core microbiome, resulting in a decrease in Oscillospiraceae abundance and a shift towards *n*-caproate production. This transition was associated with the increased dominance of other populations, including *Caproiciproducens* spp. Veillonellales-Selenomodales, and *Methanobacterium* spp. (**Fig S9**). In conclusion, these findings underscore the ecological and functional plasticity inherent to open-culture fermentations.^63,69^

In addition, our data suggests a substrate-driven ecological separation within the core microbiome. Specifically, Oscillospiraceae populations exhibited a positive correlation with high ethanol-to-lactate ratios (E-L ratio of 3), and thereby facilitating *n*-caprylate production (eigenvectors 3 and 4 in **Fig. 3A**). Conversely, *Caproiciproducens* spp. exhibited a positive correlation with lower ethanol-to-lactate ratios of 2, leading to *n*-caproate production (eigenvector 1 in **Fig. 3A**). These observations infer that Oscillospiraceae populations thrive at high ethanol concentrations, while they restrict *Caproiciproducens* spp.. Indeed, other studies have identified *Caproiciproducens* spp. to require lactate or sugars for *n*-caproate production.^80,81^ Thus, the presence of both populations in the core microbiome supports our earlier observation that ethanol and lactate as co-substrates were degraded simultaneously for chain elongation.

### 3.3. Microbiome associated with high *n*-caprylate specificity

Our findings revealed that dormant microbial populations were activated and gained abundance within the community during shifts in ethanol-to-lactate ratios and temperature. These populations, acting as specialists, worked together with the core microbiome under specific conditions, such as high ethanol-to-lactate ratios, to produce mostly *n*-caprylate (**Fig. 2A**). To explore the ecology of this correlation, we analyzed the data points separately for periods with different ethanol-to-lactate ratios (Period I-IV in **Fig. 3A**) and temperature (Period VII-XI in **Fig. 3B**), with their controls (Periods **a** and **b** for ratios and Period **d** for temperature in **Fig. S11**). During periods of ethanol-to-lactate fluctuations (Period I-IV), the microbiome shifted into two distinct clusters that correlated with product specificity (**Fig. 3A**). Notably, *Sphaerochaeta* spp., which is a key member of the core microbiome, was strongly associated with *n*-caprylate production during Periods III (**Fig. 3A**). Additionally, we observed the increased significance of other microbial populations that are not in the core microbiome. Particularly during Period III, the dbRDA analysis demonstrated a strong correlation between the populations of Erysipelaclostridiaceae UCG-004 (PERMANOVA: p=0.050; R= 0.9859), Oscillospiraceae NK4A214 (p=0.03611; R=0.9267), and *Pseudoclavibacter* spp. (p=0.00556; R=0.9635) with *n*-caprylate specificity (eigenvectors 7, 8, 11, 10, and 9 in **Fig. 3A**).

Although little is known about the Erysipelaclostridiaceae family, Palomo-Briones *et al*.^82^ observed a strong correlation between its abundance and *n*-caprylate production. Additionally, members of the Oscillospiraceae family, such as *Caproiciproducens* spp. and *Ruminococcus* spp., are known chain elongators capable of producing *n*-butyrate.^71^ Notably, our results suggest a possible interaction between Oscillospiraceae and *Pseudoclavibacter* spp. (**Fig. 3**), which is a strict aerobe. *Pseudoclavibacter* spp. has been found in various chain-elongation microbiomes, constituting approximately 30% of the total composition.^83,84^ We have recently found the importance of unplanned intrusion of oxygen into a *n*-caprylate producing microbiome and have hypothesized that *Pseudoclavibacter* spp., among aerobic yeast, removes this oxygen to convert ethanol into an intermediate that is used for chain elongation towards *n*-caprylate.^85^

During temperature fluctuations (Periods VII to XI), ecological succession occurred and influenced the structure of the community. At the lower temperatures of 25–30°C (Periods VII-VIII in **Fig. 1A**), *n*-caprylate production was superior compared to the higher temperatures of 37°C and 42°C (Period IX and X in **Fig. 1A**). We found during the temperature increase from 25°C to 42°C and then back to 30°C, that populations of *Propionibacterium* spp., *Anaerovorax* spp., Oscillospirales family, and Erysipelaclostridiaceae UCG-004 were positively correlated to *n*-caprylate production (eigenvectors 11, 10, 9, and 8, respectively, in **Fig. 3B**). However, from the literature we know that *Propionibacterium* spp. and *Anaerovorax* spp. are SCC-producing bacteria, and not chain elongators. Thus, cooperative behavior may have helped to maintain the main ecological function of *n*-caprylate production. During the increase in temperature, members of these four populations reduced in abundance, strengthening members of *Sphaerochaeta* spp. and *Caproiciproducens* spp. within the core microbiome (comparing the first two lines for both **Fig. S9D** and **Fig. S9C**), while losing *n*-caprylate production.

## ASSOCIATED CONTENT

### Supporting Information

Descriptions of bioreactor setup; calculations of production rate, specificity, and thermodynamics; additional information on bioreactor performance and microbial analysis.

## AUTHOR INFORMATION

### Author Contributions

HW performed experiments and operated the bioreactors, HW and BSJ designed the bioreactor setup, AOE and HW performed the microbiome analysis, LTA managed the project and LTA and HW designed the experiments. The manuscript was written by HW, AOE, and LTA, while all authors made edits. All authors have given approval to the final version of the manuscript.

### Notes

The authors declare no competing financial interest.

## ACKNOWLEDGMENT

HW was generously supported by the China Scholarship Council (CSC) to study at the University of Tübingen. The authors acknowledge support from the Alexander von Humboldt Foundation in the framework of the Alexander von Humboldt Professorship that is endowed by the Federal Ministry of Education and Research in Germany (to LTA.). This work or part of this work was supported by the Novo Nordisk Foundation CO_2_ Research Center (CORC) with grant number NNF21SA0072700 and is published under the number CORC_24_## to LTA. Finally, this work is also supported by the DFG through the Leibniz Prize to LTA. We further acknowledge support by the Max Planck Society to LTA as part of being a Max Planck Fellow. The authors thank Dr. Heike Budde and the Genome Center at the Max Planck Institute (MPI) for Biology Tübingen for performing the MiSeq Illumina sequencing.

