## Supplementary Information for "Steering the chain-elongating microbiome to specific medium-chain carboxylates with ethanol and lactate as co-electron donors: maximizing C8 or C6"

Number of pages: 29; Number of figures: 11; Number of tables: 9

### MATERIALS AND METHODS: Bioreactor setup

A glass-jacketed 6.5-L anaerobic reactor with a working volume of 6,00 L was used. The bioreactor was operated with constant broth recirculation through an in-line, membrane-based liquid-liquid extraction (*i.e.,* pertraction) system (**Fig. S1**). We set up two bioreactors (reactor one: R1; reactor two: R2), R1 was the operating bioreactor while R2 was the control bioreactor. The temperature was controlled using a heating bath (Huber KISS E, Germany) through all the operating conditions. The pH was measured by a probe (SL 80-425pH, Xylem Analytics, Germany) that was mounted through the lid of the bioreactor. The pH was maintained at 5.5 with an automatic controller (Eutech Instruments alpha-pH560, Vernon Hills, IL, USA) and pumps system (Masterflex® L/S® Economy Fixed-Speed Drives, OU-07540-01, Cole-Parmer Instrument Company, USA) with hydrochloric acid (*e.g.,* 4 M HCl) and sodium hydroxide (*e.g.,* 5 M NaOH).

Fresh media containing ethanol and lactate was continuously fed from a refrigerated reservoir (4$^oC$) using a peristaltic pump (Masterflex L/S® Variable-Speed Digital Drive, OU-07528-10, Cole-Parmer Instrument Company, USA) with a flow rate of 0.65 mL min$^{-1}$. The effluent was continuously recycled *via* an overflow line that was connected to the pertraction system, using a peristaltic pump (Masterflex L/S® Variable-Speed Digital Drive, OU-07528-10, Cole-Parmer Instrument Company, USA), at average folw rates of 0.94 L d$^{-1}$. The produced biogas was used to mix the broth by continuously recycling it bottom-to-top in the bioreactor. The biogas volumetric production was measured with a flow meter (BPC® µFlow, BPC Instruments, Sweden) that was connected to the biogas outlet. The biogas collection system consists of: **1)** a gas-sample septum; **2)** a glass airlock; and **3)** a two-bottle system with water as an equalization system to prevent air intrusion during sampling. Finally, a ampling port on the bioreactor was placed to take biomass samples periodically. The pertraction system was composed of two membrane contactors in series (BET area:1.4 m$^2$ each, Membrana Liqui-Cel 2.5-8, X50 membrane, Charlotte, NC, USA) that was used as the forward and backward extraction units of the MCC recovery unit (**Fig. S1**). The MCC recovery unit consisted of a pertraction using mineral oil and 30 g L$^{-1}$ tri-n-octyl phosphine oxide (TOPO) as a solvent mix (Sigma Aldrich, St. Louis, MO, USA). The extraction solution was initially buffered with 0.3 M sodium borate and then maintained at a pH=9.4 with 5 M NaOH using an automatic pump controller (Eutech Instruments alpha-pH800, the Bluelab pH Controller Connect M, Bluelab, USA). The regeneration of the extraction system was performed every eight months; membrane contactors were washed using 2% NaOH, 1% HCl, and distilled water. The extraction solution was entirely replaced to prevent losses in the extraction by solvent saturation.

### MATERIALS AND METHODS: Liquid and Gas Analysis

The concentration of carboxylates were measured by gas chromatography (GC, 7890B GC System, Agilent, USA), which was equipped with a thermal conductivity detector (TCD), using a capillary column Nukol Capillary Column (15m X 0.25 mm I.D. X 0.25um). The method was modified according to previous research[1], which was with a temperature of injection at 200°C and the detector to 250°C, ramp temperature program (initial temperature 80°C for 0.5 min, temperature ramp 20°C per 1 min to 180°C, and final temperature 180°C for 2 min), and a hydrogen flow of 21.4 mL min$^{-1}$ as a carrier gas. We measured the ethanol and lactate concentrations using a high-performance liquid chromatography (HPLC) system (Shimadzu LC 20AD), which was coupled with a refractive index and UV detector (Shimadzu, Kyoto, Japan). Separation conditions were 60$^{o}$C with 5 mM sulfuric acid as the mobile phase at a flow rate of 0.6 mL min$^{-1}$ in an Aminex HPX-87H column (Bio-Rad, Hercules, CA, USA). *Prior* to the analysis, samples were filtered through a sterile Acrodisc 0.22-mm pore size, polyvinylidene fluoride membrane syringe filter (Pall Life Sciences, Port Washington, NY, USA) to remove possible biological and particulate contaminants. Finally, production rates for MCCs (mmol C L$^{-1}$ d$^{-1}$) were calculated using the MCC concentrations in the pertraction system and the effluent of the bioreactor (**Table S5**), according to Xu *et al.* [2]

### MATERIALS AND METHODS: Calculations

#### 1. The Equation of Volumetric Production Rate on Day n (mmol C $L^{-1}$ $d^{-1}$)

Equation $\frac{C_{e,n}V}{HRT} + \frac{(C_{b,n} - C_{b,n-1})V_b}{T_n - T_{n-1}} \quad \frac{1}{1000V}$

| | |
|---|---|
| $C_{e,n}$ | concentration of carboxylates in effluent on the day n, mM |
| V | volume of reactor, L |
| HRT | hydraulic retention time on the day n, d |
| $C_{b,n} - C_{b,n-1}$ | concentrations of carboxylates in the stripping solution on the day n and n-1, mM |
| $V_b$ | volume of the stripping solution on the day n, L |
| $T_n$ | the day n, d |

#### 2. Specificities

The specificities (%) were calculated based on the specific carboxylate product rate in mmol C $L^{-1}$ $d^{-1}$ divided by to the production rate for the sum of all produced carboxylates in mmol C $L^{-1}$ $d^{-1}$.

#### 3. Thermodynamics of Biochemical Reactions

The thermodynamic calculation of biochemical reactions was based on the study by Alberty *et al.*[3, 4].

The transformed Gibbs free energy ($\Delta rG_T^{'}$):

$$\Delta rG_T^{'} = \Delta rG_T^{'0} + RT\ln Q \qquad \textbf{Eq. S1}$$

where $\Delta rG_T^{'0}$ is the standard transformed Gibbs free energy of a reaction at a temperature (T). Q is a factor related to the activities of reactants and products defined as:

$$Q = \frac{(\alpha_A)^a(\alpha_B)^b\ldots(\alpha_C)^c}{(\alpha_X)^x(\alpha_Y)^y\ldots(\alpha_Z)^z} \qquad \textbf{Eq. S2}$$

where the numerator represents the activity of products A, B, C, *etc*. and the denominator represents the activity of reactants X, Y, Z, *etc*. The powers are the stoichiometric coefficients of the products and reactants in each reaction.

To obtain $\Delta rG_T^{'0}$, the standard Gibbs free energy of formation ($\Delta_f G^0$) and standard enthalpy of formation ($\Delta_f H^0$) for each reactant and product at T = 298.15 K and ionic strength (I) = 0 M were found in a reference[5]. For those chemicals for which values of standard Gibbs free energy of formation ($\Delta_f G^0$) and standard enthalpy of formation ($\Delta_f H^0$) were not available, we calculated those values based on the methods from Mavrovouniotis[6] and Hanselmann[7].

For a given condition of 303 K, the Gibbs free energy of formation at T = 303 K, $\Delta_f G^0_{i,303}$ is adjusted with **Eq. S3:**

$$\Delta_f G^0_{i,303k} = (\frac{303k}{298.15k}) \times \Delta_f G^0_{i,298.15k} + (1 - \frac{303k}{298.15k}) \times \Delta_f H^0_{i,298.15} \qquad \textbf{Eq. S3}$$

Subsequently, the standard transformed Gibbs free energies of formation at different pH and ionic strength, $\Delta_f G^0_{i,310k}{}^{(pH, I)}$ are calculated as follows:

$$\Delta_f G'^0_{I,303k}(pH, I) = \Delta_f G^0_{i,303k} - N_{H,i} RT\ln 10^{-pH} - RT\alpha(Z_i^2 - N_{H,i}) I^{1/2}/(1 + BI^{1/2}) \qquad \textbf{Eq. S4}$$

where $RT\alpha = 9.20483 \times 10^{-3}\ T - 1.28467 \times 10^{-5} T^2 + 4.95199 \times 10^{-8} T^3$, $B = 1.6\ kg^{1/2} mol^{-1/2}$, $N_{H,i}$ is the number of hydrogen atoms in a substance, and $Z_i$ is the charge number. Finally, with the standard transformed Gibbs free energies of formation of each reactant and product, the standard transformed Gibbs free energy of each biochemical reaction is calculated using **Eq. S5**,

$$\Delta_r G'^0_T = \sum Vprod \Delta_f G'^0_T - \sum VReac \Delta_f G'^0_T \qquad \textbf{Eq. S5}$$

Here, we evaluated two conditions for thermodynamic analysis; one was the standard condition, and another one was with a pH at 5.5 and a temperature of 30°C.

**FIGURES:**

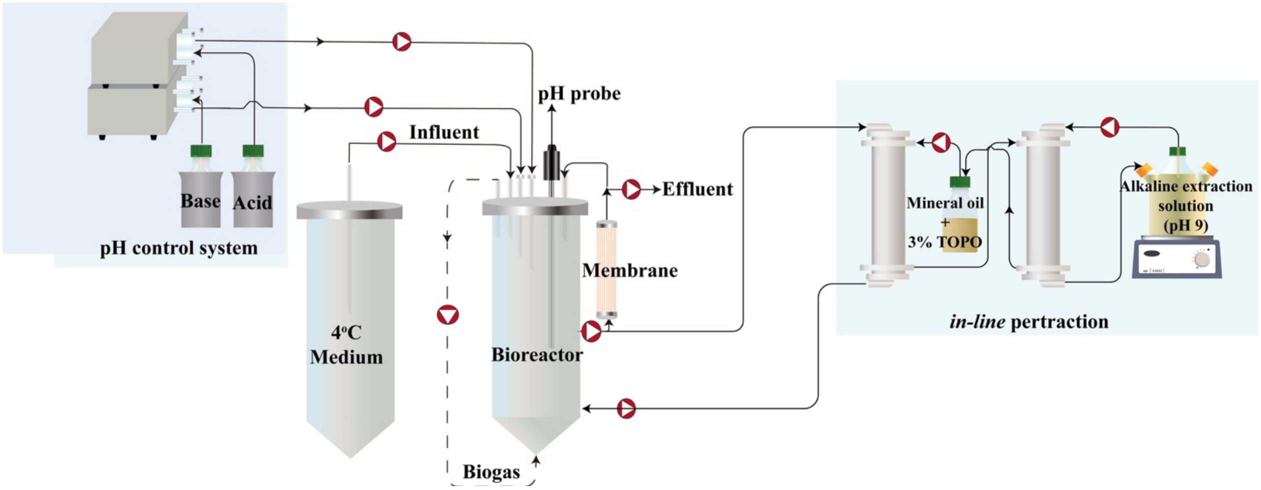


**Fig. S1.** Set up of the reactor system.

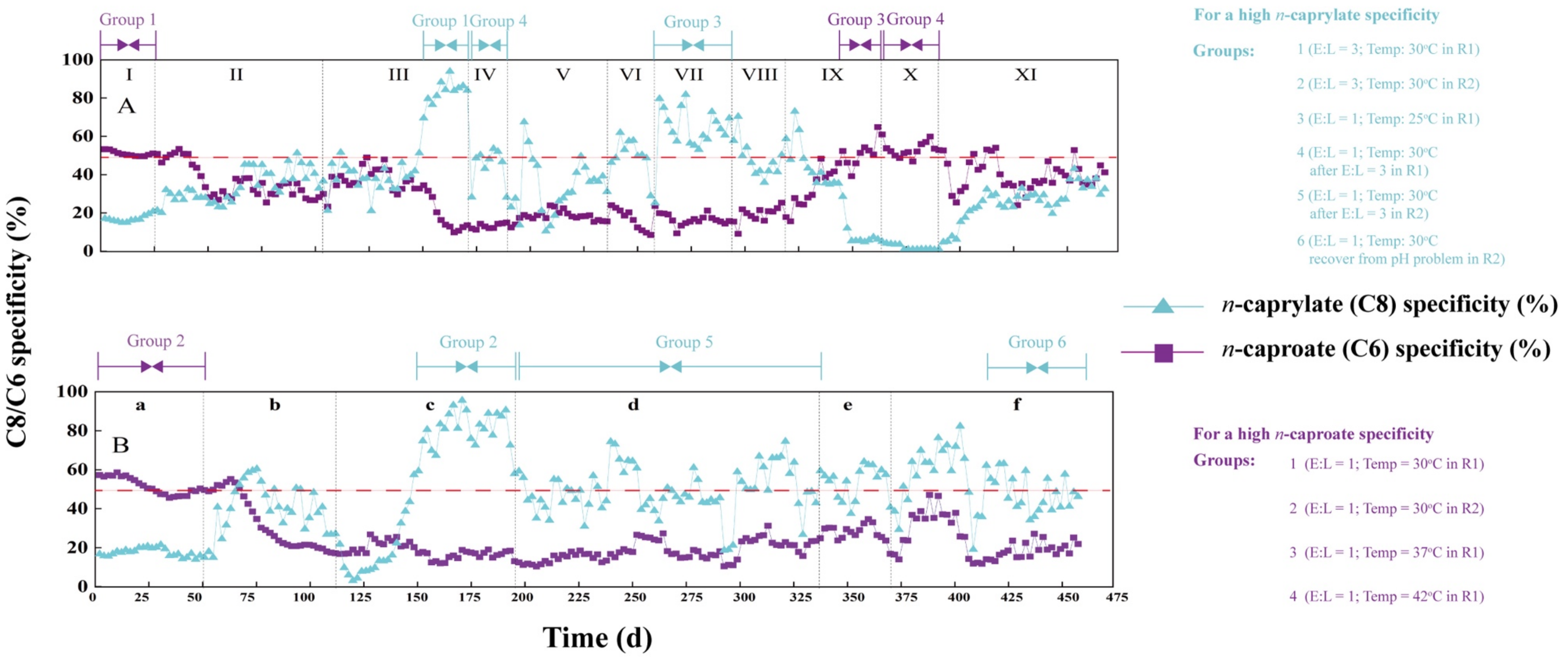


**Fig. S2.** The specificity of *n*-caprylate or *n*-caproate during the operating period for: (A) R1; and (B) R2. We observed six and four groups that clustered, respectively. For a high *n*-caprylate specificity, the six groups consisted of three different operating conditions (E-L ratio = 3 *plus* temperature = 30ºC; E-L ratio = 1 *plus* temperature = 25ºC; E-L ratio = 1 *plus* temperature = 30ºC but through hysteresis after a high *n*-caprylate specificity period; and). For a high *n*-caproate specificity, the four groups consisted of three different operating conditions (E-L ratio = 1 *plus* temperature = 30ºC; E-L ratio = 1 *plus* temperature = 37ºC; and E-L ratio = 1 *plus* temperature = 42ºC).

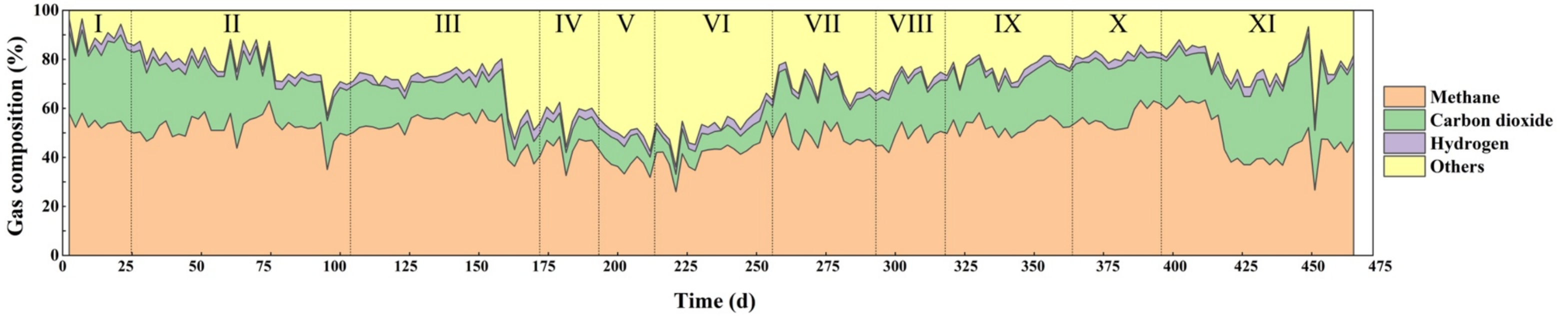


**Fig. S3.** Biogas composition throughout the operating periods for the bioreactor (R1). Methane, carbon dioxide, and hydrogen percentages in the biogas.

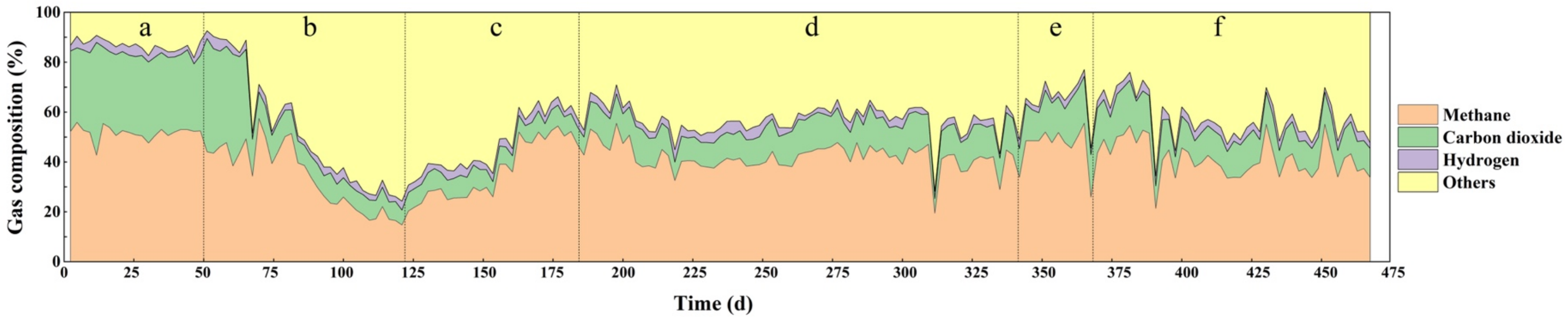


**Fig. S4.** Biogas composition throughout the operating periods for the bioreactor (R2). Methane, carbon dioxide, and hydrogen percentages in the biogas.

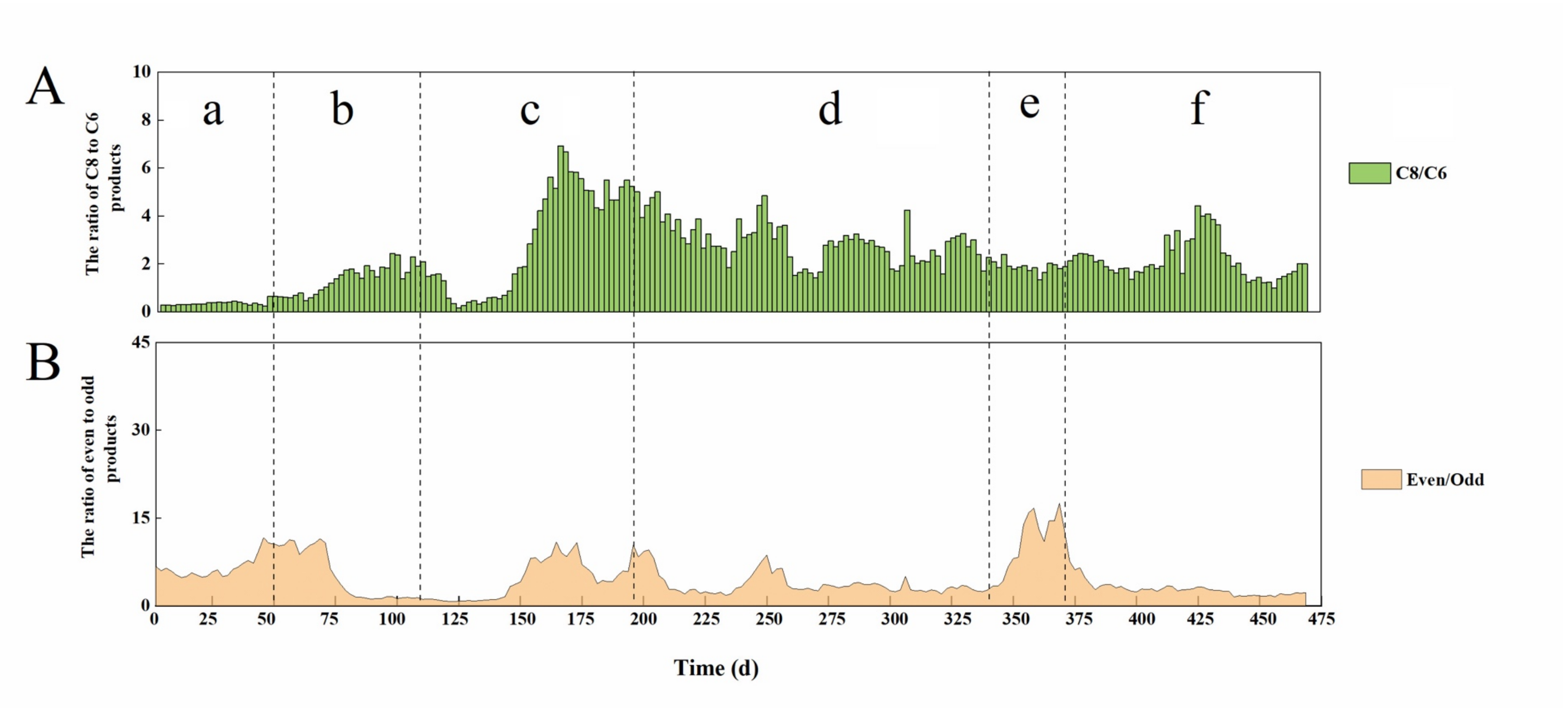


**Fig. S5.** Performance for the control bioreactor (R2) during Periods **a-f**: (A) column chart for ratio of C8 to C6; (B) stacked chart for ratio of even to odd. The data represent a 6-day moving average.

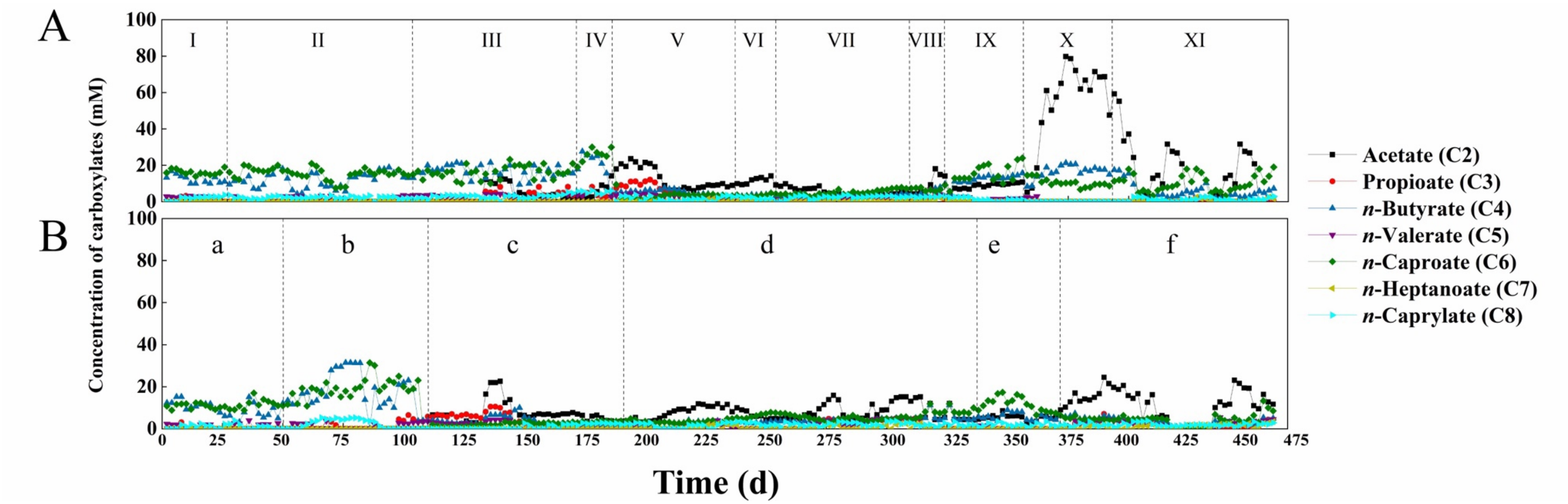


**Fig. S6.** The concentrations of carboxylates in the bioreactor: (A) R1; and (B) R2.

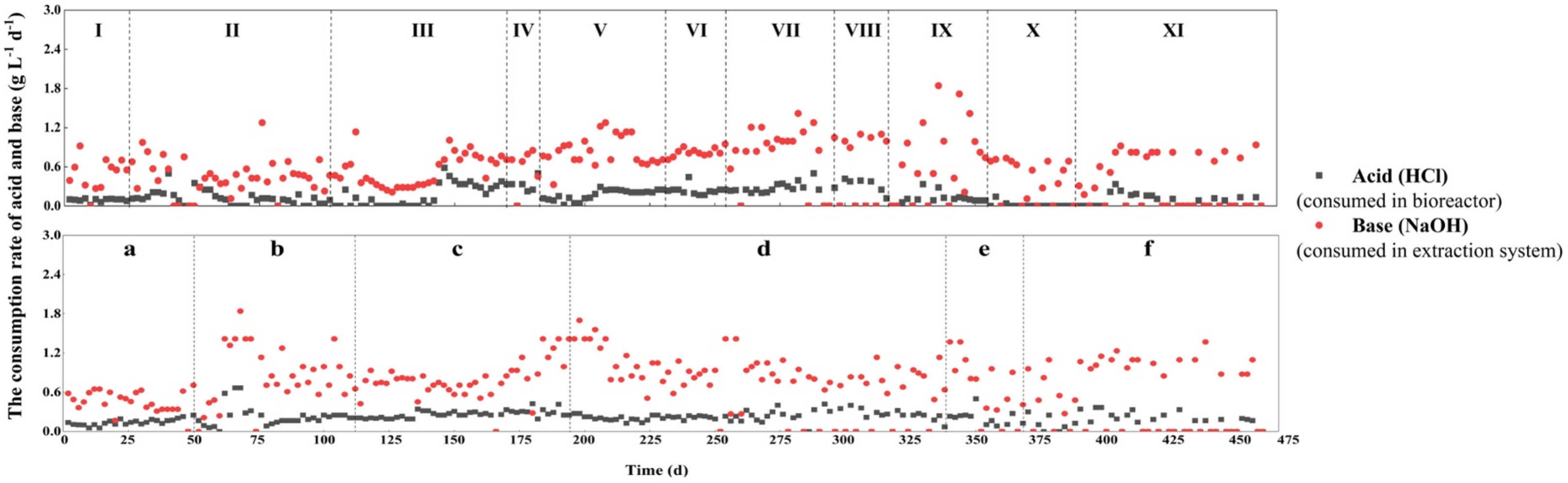


**Fig. S7.** The acid and base consumption rate for: (A) R1; and (B) R2.

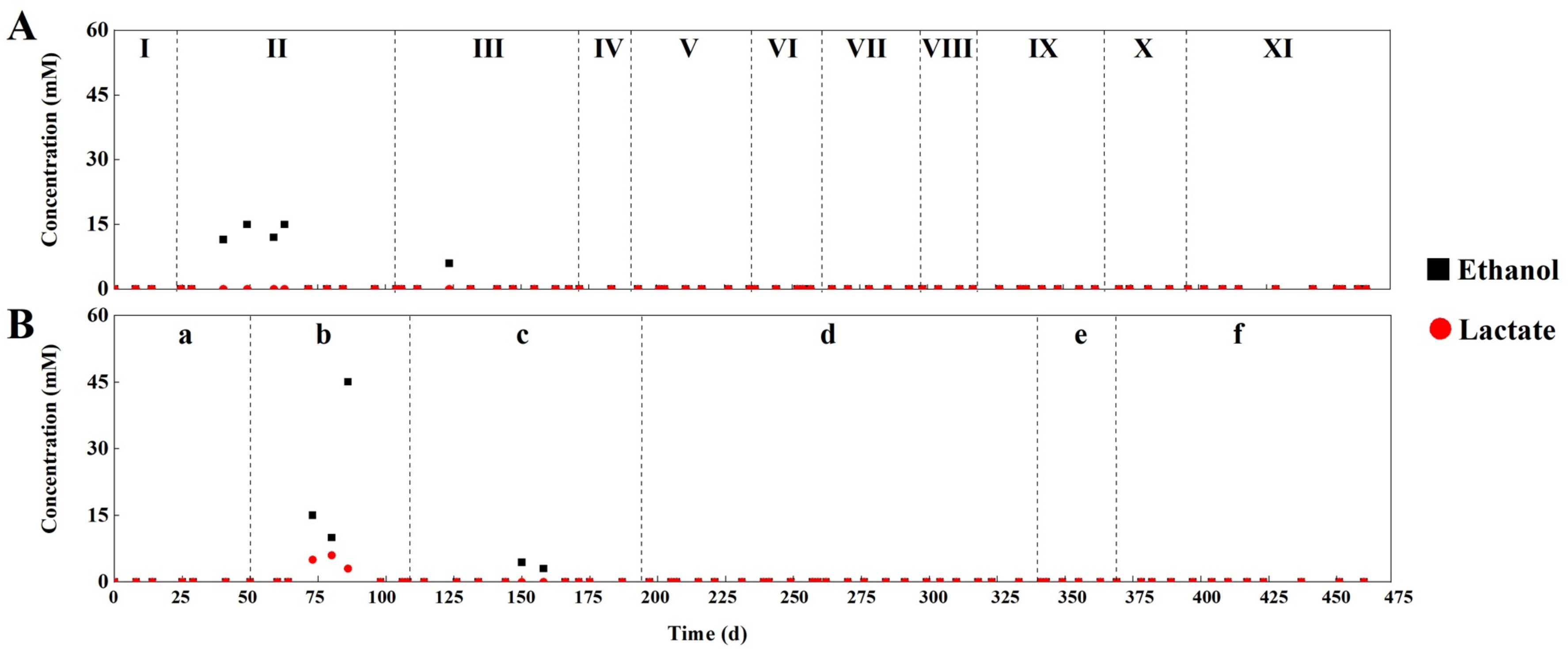


**Fig. S8.** The concentrations of ethanol and lactate in the bioreactor: (A) R1; and (B) R2.

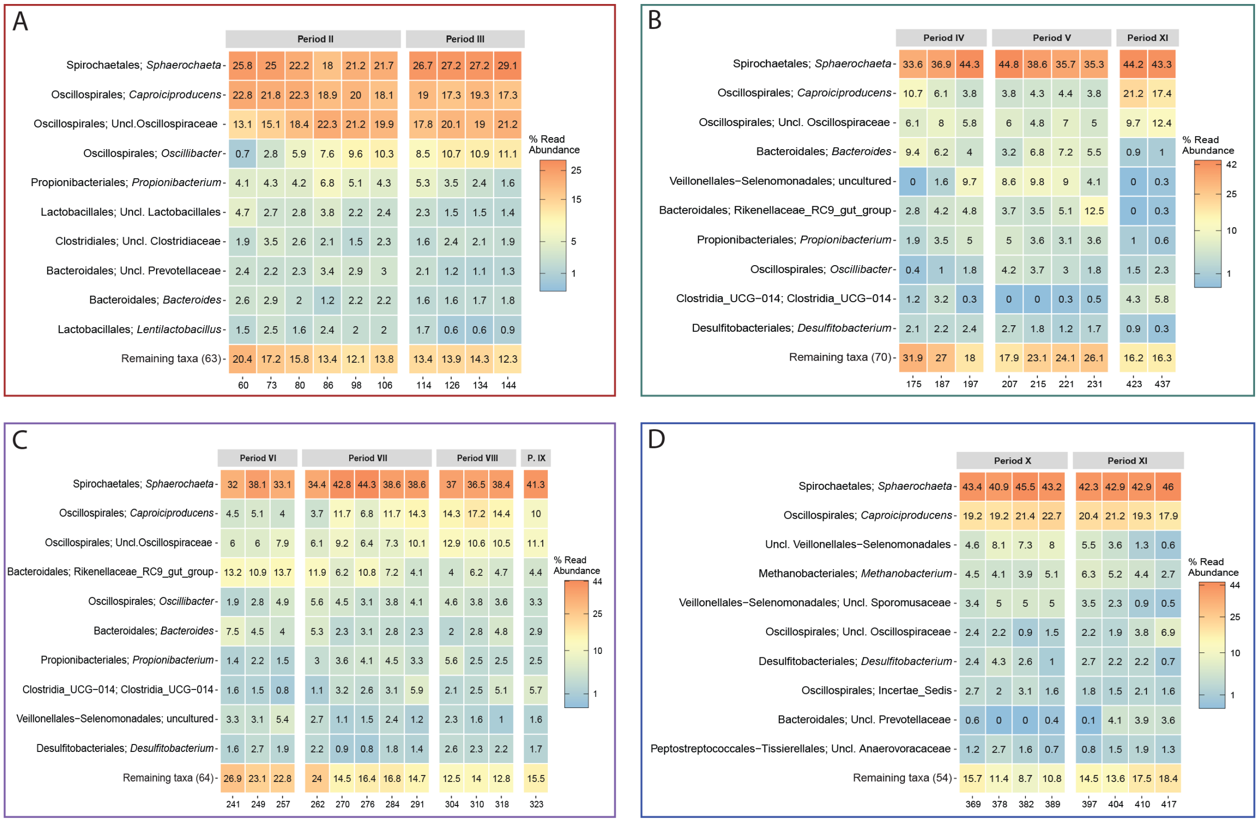


**Fig. S9.** Heatmaps of the microbial composition within the clusters identified in the experimental bioreactor (R1) with dbRDA (**Fig. 2A**). Relative abundances were normalized by the read number *per* sample. Heatmaps show the 10 more abundant taxa on each cluster, while the X axis represents the day of sampling.

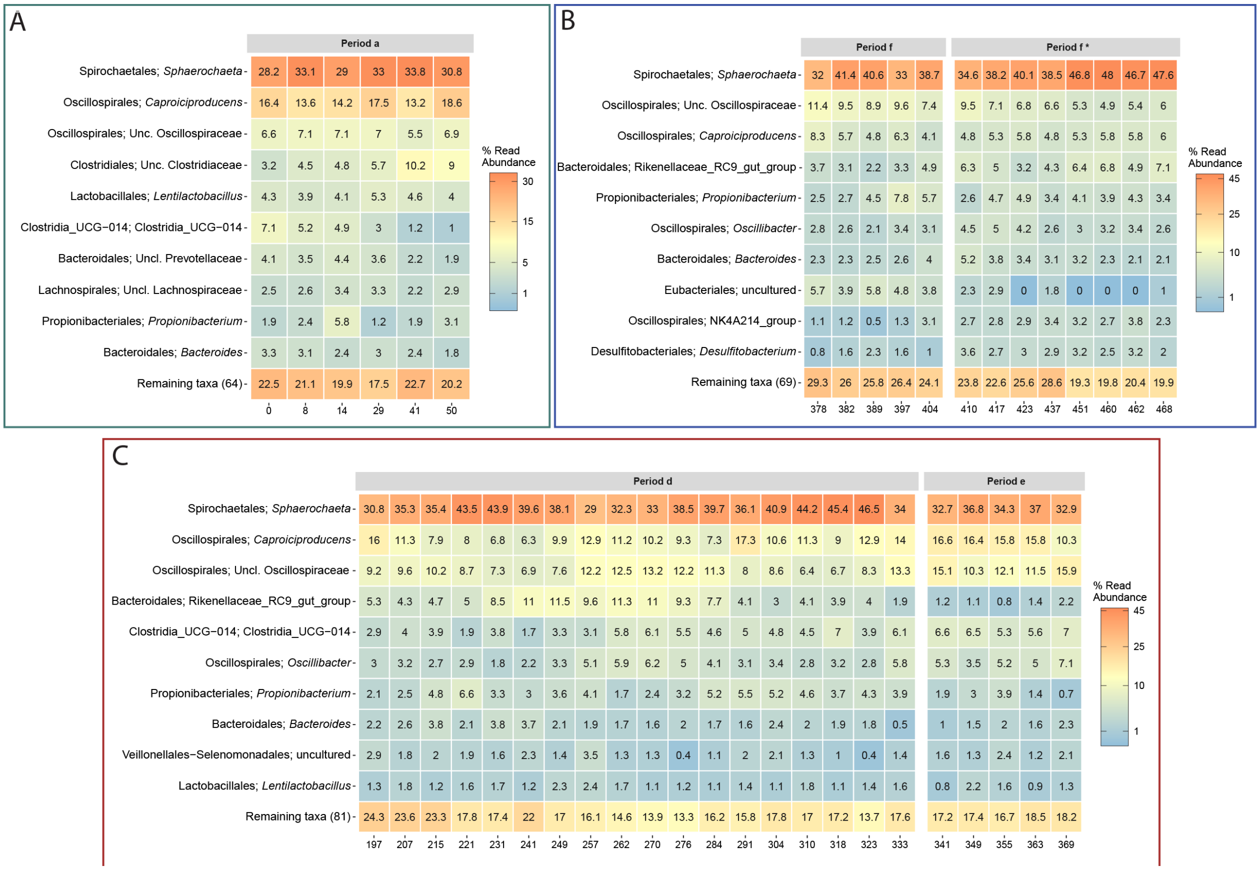


**Fig. S10.** Heatmaps of the microbial composition within the clusters identified in the control bioreactor (R2) with dbRDA (**Fig. 2B**). Relative abundances were normalized by the reads number *per* sample, heatmaps show the 10 more abundant taxa on each cluster, X axis represents the day of sampling.

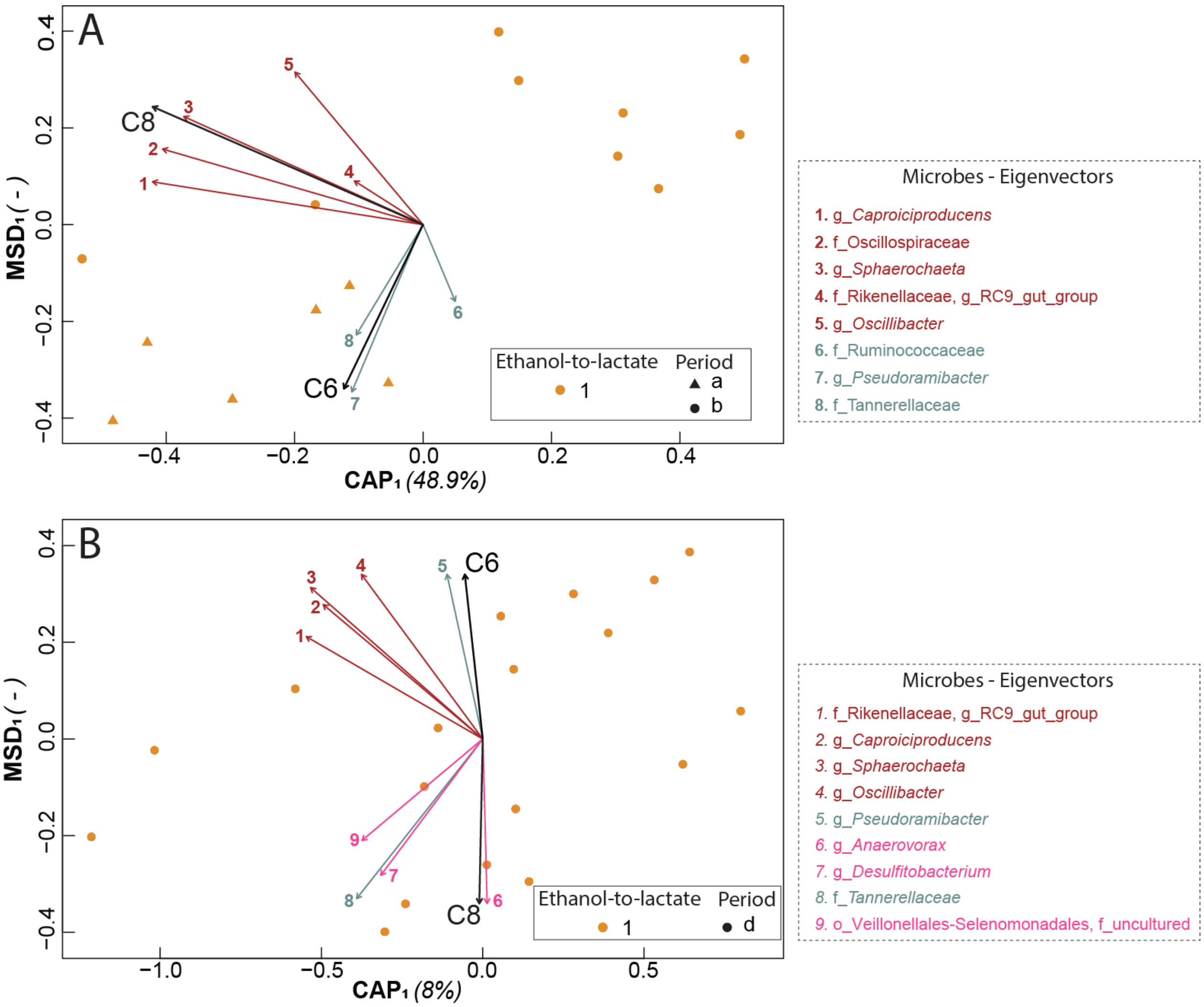


**Fig. S11.** Component analyses of control Periods **a** and **b** for ethanol-to-lactate ratios in R2(**A**) and control Period **d** for temperature variations in R2 (**B**) using distance-based redundancy analyses with Bray-Curtis ß-diversity distance matrixes. Eigenvectors where calculated based on PERMANOVA significances < 0.05, Eigenvectors: core microbiome members (Brown); microbes with low significance (Blue); significant microbes with correlation coefficients <0.08 (Pink); highly significant microbes with correlation coefficients >0.08 (Green). Other vectors: C6 (*n*-caproate), C8 (*n*-caprylate).

### TABLES

**Table S1.** The components of the basal medium.

| Components [a] | $L^{-1}$ |
|---|---|
| MQ water | 772 mL |
| Mineral stock solution (ml, 10×) | 100 ml |
| $KH_2PO_4$ | 0.23 g |
| Yeast extract | 1.25 g |
| $Na_2CO_3$ | 3 g |
| L-cystein HCl (ml,100×) | 10 mL |
| Vitamin solution (ml, 100×) | 5 mL |

[a] The ethanol and lactate were added differently according to the experiment design.

**Table S2**. The components of mineral stock solution (10×).

| Components | g $L^{-1}$ |
|---|---|
| NaCl | 11.7 |
| $NH_4Cl$ | 2.37 |
| $CaCl_2$•$2H_2O$ | 0.65 |
| $MgCl_2$•$6H_2O$ | 0.25 |
| $MnCl_2$•$4H_2O$ | 0.31 |
| $ZnCl_2$•$H_2O$ | 0.12 |
| $CoCl_2$•$6H_2O$ | 0.048 |

**Table S3.** The compounds of vitamin solution (100×).

| Components | mg $L^{-1}$ |
|---|---|
| Pyridoxine | 20 |
| Thiamine | 10 |
| Riboflavin | 10 |
| Calcium pantothenate | 10 |
| Thioctic acid | 10 |
| Para-aminobenzoic acid | 10 |
| Nicotinic acid | 10 |
| Vitamin $B_{12}$ | 10 |
| d-Biotin | 4 |
| Folic acid | 4 |
| 2-Mercaptoethanesulfonic acid | 4 |

**Table S4.** The operating conditions for different periods for R1and R2.

| | | E-L ratio | Temperature (°C) | pH | HRT (d) | OLR (mmol C $L^{-1}$ $d^{-1}$) | Time (d) | Periods of steady-state conditions (d) |
|---|---|---|---|---|---|---|---|---|
| | | | | **R1** | | | | |
| Period I | | 1 | 30 | 5.5 | 6.4 | 110 | 1-25 | 1-25 |
| Period II | | 2 | 30 | 5.5 | 6.4 | 110 | 26-107 | 65-107 |
| Period III | | 3 | 30 | 5.5 | 6.4 | 110 | 108-171 | 158-171 |
| Period IV | | 1 | 30 | 5.5 | 6.4 | 110 | 172-187 | 172-187 |
| Period V (dysfunctional extraction) | | 3 | 30 | 5.5 | 6.4 | 110 | 188-239 | |
| Period VI (recovery) | | 1 | 30 | 5.5 | 6.4 | 110 | 240-259 | 240-259 |
| Period VII | | 1 | 25 | 5.5 | 6.4 | 110 | 260-299 | 260-299 |
| Period VIII | | 1 | 30 | 5.5 | 6.4 | 110 | 300-318 | 300-318 |
| Period IX | | 1 | 37 | 5.5 | 6.4 | 110 | 319-363 | 334-363 |
| Period X | | 1 | 42 | 5.5 | 6.4 | 110 | 364-395 | 381-395 |
| Period XI | | 1 | 30 | 5.5 | 6.4 | 110 | 396-468 | 418-468 |
| | | | | **R2** | | | | |
| Period a (control for Periods I-IV) | | 1 | 30 | 5.5 | 6.4 | 110 | 0-50 | 0-50 |
| Period b | (improving pump rates) | 1 | 30 | 5.5 | 6.4 | 110 | 51-95 | |
| | (dysfunctional extraction) | 1 | 30 | 5.5 | 6.4 | 110 | 96-119 | |
| Period c (recovery) | | 1 | 30 | 5.5 | 6.4 | 110 | 120-149 | |
| | | 3 | 30 | 5.5 | 6.4 | 110 | 150-190 | |
| Period d (control for Periods VII-XI) | | 1 | 30 | 5.5 | 6.4 | 110 | 191-339 | 198-339 |
| Period e (failure of pH probe) | | 1 | 30 | 5.5 | 6.4 | 110 | 340-371 | |
| Period f (recovery) | | 1 | 30 | 5.5 | 6.4 | 110 | 372-471 | 372-471 |

*E: ethanol; and L: L-lactate.

**Table S5.** The average volumetric MCC production rates (mmol C $L^{-1}$ $d^{-1}$) and ratios (mol C mol $C^{-1}$) during different operating conditions for R1 and R2.

| **Operating condition** | | Periods | Acetate | Propionate | *n*-Butyrate | *n*-Valerate | *n*-Caproate | *n*-Heptanoate | *n*-Caprylate | *n*-Nonanoate | C8/C6 | Even/Odd |
|---|---|---|---|---|---|---|---|---|---|---|---|---|
| | | | | | | **R1** | | | | | | |
| E-L ratio | 1 | I | 0 | 0 | 9.90±0.17 | 3.38±0.45 | 47.0±0.16 | 10.9±0.20 | 16.0±1.97 | 0.92±0.18 | 0.34±0.04 | 4.90±0.70 |
| | 2 (non-steady state) | II | 0 | 0 | 8.27±0.22 | 2.53±0.38 | 37.1±1.04 | 6.65±0.43 | 25.4±1.09 | 0.53±0.19 | 0.71±0.15 | 7.48±1.76 |
| | 2 | | 0 | 0 | 10.2±0.85 | 2.36±0.51 | 29.4±1.37 | 6.61±0.38 | 37.2±2.45 | 1.19±0.31 | 1.28±0.21 | 7.77±1.58 |
| | 3 (non-steady state) | III | 0 | 0 | 12.9±0.64 | 3.08±0.71 | 33.8±1.67 | 5.72±0.24 | 39.7±0.15 | 1.81±0.64 | 1.22±0.51 | 8.31±1.59 |
| | 3 | | 0 | 0 | 2.17±0.53 | 0.93±0.12 | 11.02±1.05 | 4.40±0.79 | 79.3±1.55 | 1.94±0.57 | 7.56±1.46 | 13.3±2.37 |
| | 1 | IV | 0 | 0 | 1.45±0.15 | 2.61±0.14 | 11.9±1.22 | 9.89±0.31 | 42.8±0.37 | 3.88±0.27 | 3.61±0.67 | 3.73±1.09 |
| Dysfunctional extraction system | | V | 2.81±0.18 | 1.98±0.82 | 5.23±0.21 | 7.75±0.32 | 16.7±0.56 | 13.8±0.38 | 30.9±2.95 | 5.50±1.45 | 1.91±0.81 | 1.94±0.40 |
| Recovery period | | VI | 3.45±0.64 | 0 | 3.86±0.90 | 4.64±0.30 | 15.2±1.10 | 10.7±0.11 | 46.5±0.32 | 3.26±1.45 | 3.34±1.01 | 4.11±1.27 |
| Operating temperature (ºC) | 25 (non-steady state) | VII | 2.59±0.15 | 0 | 4.86±0.71 | 2.66±0.71 | 18.2±1.33 | 9.46±0.45 | 57.2±0.97 | 6.32±0.93 | 3.26±1.24 | 4.64±1.37 |
| | 25 | | 1.58±0.52 | 0.23±0.01 | 2.68±0.65 | 2.45±0.53 | 14.0±0.78 | 8.36±0.38 | 59.2±0.86 | 3.04±0.15 | 4.45±1.32 | 5.61±0.97 |
| | 30 | VIII | 1.38±0.12 | 0.39±0.25 | 2.60±0.18 | 1.36±0.17 | 18.8±1.55 | 9.10±0.41 | 41.4±1.52 | 1.72±0.81 | 2.22±0.28 | 5.66±2.23 |
| | 37 (non-steady state) | IX | 3.14±0.23 | 0.66±0.16 | 5.49±0.65 | 6.55±0.21 | 21.3±1.79 | 12.2±0.23 | 51.5±1.04 | 4.03±0.60 | 2.52±0.66 | 3.55±0.73 |
| | 37 | | 2.85±0.28 | 0 | 9.54±0.61 | 2.76±0.27 | 41.5±1.32 | 6.45±0.16 | 20.6±1.44 | 2.01±0.46 | 0.54±0.43 | 13.6±0.34 |
| | 42 (non-steady state) | X | 2.85±0.28 | 0 | 11.4±0.50 | 2.42±0.38 | 55.2±2.08 | 1.26±0.26 | 4.73±0.96 | 0.09±0.02 | 0.09±0.01 | 20.6±2.62 |
| | 42 | | 18.5±0.33 | 0 | 15.8±0.32 | 1.23±0.29 | 48.1±1.49 | 1.04±0.28 | 0.89±0.49 | 0.56±0.54 | 0.04±0.03 | 30.6±2.13 |
| | 30 (non-steady state) | XI | 11.1±0.51 | 0 | 12.2±0.61 | 1.30±0.63 | 37.7±1.02 | 2.97±0.73 | 18.6±0.87 | 0.79±0.52 | 0.47±0.13 | 16.1±0.90 |
| | 30 | | 0.74±0.32 | 0 | 4.22±0.46 | 3.13±0.39 | 35.4±1.51 | 6.30±1.44 | 27.5±0.24 | 0.03±0.03 | 0.80±0.18 | 7.23±1.05 |

| | | | | | | | | | | | |
|---|---|---|---|---|---|---|---|---|---|---|---|
| | | | | | **R2** | | | | | | |
| Control of the E-L ratio experiment | a | 0 | 0.07±0.01 | 7.80±0.97 | 2.39±0.73 | 42.5±1.64 | 7.48±1.09 | 15.9±0.41 | 0.66±0.37 | 0.38±0.11 | 6.69±1.98 |
| Improving pump rate | b | 0 | 0.95±0.17 | 5.84±0.17 | 2.70±0.66 | 34.9±2.07 | 10.3±0.94 | 36.4±0.97 | 4.45±0.07 | 1.15±0.49 | 7.13±2.07 |
| Dysfunctional extraction system | b | 0.37±0.13 | 4.28±0.73 | 5.44±0.79 | 8.74±0.75 | 17.1±0.97 | 23.4±1.82 | 32.5±1.60 | 5.03±0.21 | 1.89±0.38 | 1.33±0.16 |
| Recovery period | c | 2.58±0.70 | 6.17±0.49 | 8.98±0.37 | 13.2±0.37 | 16.3±1.88 | 17.4±1.77 | 10.2±0.65 | 4.78±0.81 | 0.65±0.47 | 0.92±0.14 |
| | c | 2.05±0.52 | 0.49±0.11 | 3.55±0.84 | 2.84±0.90 | 14.4±1.04 | 7.84±0.01 | 55.7±1.94 | 3.35±0.05 | 4.21±1.83 | 6.37±2.80 |
| Control of the operating temperature experiment | d | 2.62±0.78 | 0.66±0.19 | 2.95±0.05 | 2.54±0.44 | 14.7±1.92 | 10.5±0.45 | 43.1±1.27 | 5.11±0.47 | 3.07±1.00 | 3.89±1.98 |
| Dysfunctional pH sensor | e | 1.63±0.21 | 0 | 4.42±0.46 | 1.22±0.41 | 22.9±1.24 | 6.72±0.36 | 43.2±0.79 | 1.72±0.83 | 1.90±0.24 | 10.4±5.16 |
| Recovery period | f | 3.44±0.90 | 0.69±0.21 | 2.81±0.58 | 4.03±0.09 | 20.4±1.52 | 15.8±0.15 | 40.9±1.29 | 5.17±0.43 | 2.16±0.83 | 2.87±1.24 |

**Table S6.** The operating conditions and selectivity and specificity during different operating conditions for R1 and R2.

| **Operating condition** | | | Volumetric ethanol loading rate (mmol C $L^{-1}$ $d^{-1}$) | Volumetric lactate loading rate (mmol C $L^{-1}$ $d^{-1}$) | Combined volumetric ethanol and lactate loading rate (mmol C $L^{-1}$ $d^{-1}$) | Selectivity | | | Specificity | | |
|---|---|---|---|---|---|---|---|---|---|---|---|
| | | | | | | C6/E+L fed | C8/E+L fed | MCCs/E+L fed | C6/all carboxylates | C8/all carboxylates | MCCs/all carboxylates |
| | | | | | | **R1** | | | | | |
| E-L_ratio | 1 | I | 44 | 66 | 110 | 0.43±0.01 | 0.15±0.02 | 0.68±0.04 | 0.54±0.02 | 0.18±0.02 | 0.85±0.02 |
| | 2 (non-steady state) | II | 63 | 47 | | 0.34±0.08 | 0.23±0.03 | 0.63±0.10 | 0.46±0.06 | 0.32±0.03 | 0.86±0.02 |
| | 2 | | | | | 0.27±0.03 | 0.34±0.05 | 0.68±0.07 | 0.34±0.04 | 0.43±0.03 | 0.85±0.05 |
| | 3 (non-steady state) | III | 74 | 36 | | 0.31±0.05 | 0.36±0.12 | 0.74±0.13 | 0.35±0.06 | 0.40±0.09 | 0.83±0.05 |
| | 3 | | | | | 0.12±0.03 | 0.72±0.03 | 0.89±0.04 | 0.12±0.02 | 0.78±0.04 | 0.97±0.01 |
| | 1 | IV | 44 | 66 | | 0.11±0.01 | 0.39±0.07 | 0.62±0.10 | 0.17±0.03 | 0.59±0.05 | 0.95±0.03 |
| Dysfunctional extraction system | | V | | | | 0.15±0.02 | 0.28±0.12 | 0.61±0.13 | 0.20±0.04 | 0.37±0.10 | 0.36±0.10 |
| Recovery period | | VI | | | | 0.14±0.05 | 0.42±0.08 | 0.69±0.16 | 0.17±0.03 | 0.56±0.07 | 0.53±0.06 |
| Operating temperature (ºC) | 25 (non-steady state) | VII | 44 | 66 | 110 | 0.17±0.02 | 0.52±0.18 | 0.83±0.16 | 0.19±0.06 | 0.55±0.13 | 0.90±0.02 |
| | 25 | | | | | 0.13±0.03 | 0.54±0.07 | 0.77±0.08 | 0.15±0.03 | 0.65±0.04 | 0.92±0.01 |
| | 30 | VIII | | | | 0.17±0.02 | 0.38±0.05 | 0.65±0.06 | 0.25±0.03 | 0.54±0.03 | 0.93±0.02 |
| | 37 (non-steady state) | IX | | | | 0.19±0.04 | 0.47±0.09 | 0.81±0.13 | 0.20±0.05 | 0.49±0.04 | 0.85±0.02 |
| | 37 | | | | | 0.38±0.06 | 0.19±0.13 | 0.64±0.17 | 0.51±0.16 | 0.22±0.12 | 0.82±0.03 |
| | 42 (non-steady state) | X | | | | 0.5±0.05 | 0.04±0.01 | 0.56±0.06 | 0.68±0.03 | 0.06±0.01 | 0.76±0.04 |
| | 42 | | | | | 0.44±0.03 | 0.02±0.01 | 0.47±0.03 | 0.55±0.04 | 0.02±0.02 | 0.59±0.04 |

| | | | | | | | | | | |
|---|---|---|---|---|---|---|---|---|---|---|
| 30 (non-steady state) | XI | | | | 0.34±0.09 | 0.17±0.07 | 0.55±0.14 | 0.44±0.08 | 0.21±0.08 | 0.70±0.14 |
| 30 | | | | | 0.32±0.06 | 0.25±0.05 | 0.63±0.09 | 0.46±0.07 | 0.36±0.04 | 0.89±0.02 |
| | | | | | **R2** | | | | | |
| Control of the E-L ratio experiment | a | | | | 0.39±0.03 | 0.14±0.04 | 0.60±0.06 | 0.56±0.03 | 0.21±0.05 | 0.87±0.03 |
| Improving pump rate | b | 44 | 66 | | 0.32±0.08 | 0.33±0.09 | 0.78±0.09 | 0.38±0.14 | 0.38±0.06 | 0.90±0.03 |
| Dysfunctional extraction system | b | | | | 0.15±0.01 | 0.29±0.06 | 0.71±0.07 | 0.18±0.01 | 0.33±0.05 | 0.8±0.03 |
| Recovery period | c | 74 | 36 | 110 | 0.15±0.03 | 0.09±0.06 | 0.44±0.08 | 0.21±0.03 | 0.13±0.08 | 0.61±0.08 |
| | c | 44 | 66 | | 0.13±0.03 | 0.51±0.17 | 0.74±0.14 | 0.16±0.05 | 0.60±0.16 | 0.89±0.07 |
| Control of the operating temperature experiment | d | | | | 0.13±0.04 | 0.39±0.11 | 0.67±0.15 | 0.18±0.03 | 0.52±0.10 | 0.89±0.05 |
| Dysfunctional pH sensor | e | 44 | 66 | | 0.21±0.03 | 0.39±0.06 | 0.68±0.09 | 0.28±0.04 | 0.53±0.04 | 0.91±0.02 |
| Recovery period | f | | | | 0.19±0.07 | 0.37±0.12 | 0.75±0.20 | 0.22±0.04 | 0.44±0.09 | 0.88±0.04 |

**Table S7.** The average volumetric acid and base consumption rates during different operating conditions for R1 and R2.

| **Operating condition** | | Periods | Acid (HCL/ g $L^{-1}$ $d^{-1}$) | Base (NaOH/ g $L^{-1}$ $d^{-1}$) |
|---|---|---|---|---|
| R1 | | | | |
| E-L ratio | 1 | I | 0.09±0.03 | 0.5±0.24 |
| | 2 (non-steady state) | II | 0.18±0.12 | 0.41±0.3 |
| | 2 | | 0.06±0.06 | 0.46±0.26 |
| | 3 (non-steady state) | III | 0.07±0.15 | 0.43±0.21 |
| | 3 | | 0.33±0.07 | 0.76±0.15 |
| | 1 | IV | 0.26±0.15 | 0.62±0.28 |
| Dysfunctional extraction system | | V | 0.18±0.08 | 0.84±0.23 |
| Recovery period | | VI | 0.24±0.08 | 0.82±0.05 |
| Operating temperature (°C) | 25 (non-steady state) | VII | 0.19±0.11 | 0.64±0.39 |
| | 25 | | 0.25±0.13 | 0.92±0.41 |
| | 30 | VIII | 0.18±0.2 | 0.51±0.54 |
| | 37 (non-steady state) | IX | 0.09±0.09 | 0.61±0.5 |
| | 37 | | 0.11±0.09 | 0.76±0.61 |
| | 42 (non-steady state) | X | 0 | 0.47±0.4 |
| | 42 | | 0 | 0.37±0.3 |
| | 30 (non-steady state) | XI | 0.08±0.13 | 0.36±0.34 |
| | 30 | | 0.06±0.07 | 0.34±0.41 |
| R2 | | | | |
| Control of the E-L ratio experiment | | a | 0.14±0.06 | 0.46±0.17 |
| Improving pump rate | | b | 0.22±0.2 | 0.86±0.55 |
| Dysfunctional extraction system | | b | 0.22±0.03 | 0.81±0.26 |
| Recovery period | | c | 0.24±0.05 | 0.76±0.11 |
| | | c | 0.27±0.08 | 0.77±0.32 |
| Control of the operating temperature experiment | | d | 0.21±0.11 | 0.82±0.43 |
| Dysfunctional pH sensor | | e | 0.17±0.13 | 0.69±0.45 |
| Recovery period | | f | 0.11±0.13 | 0.5±0.5 |

**Table S8.** Thermodynamic calculations of the reactions (KJ $mol^{-1}$).

| Eq. | Bioprocess | Reaction [a] | 25ºC | | 30ºC | | 37ºC | | 42ºC | |
|---|---|---|---|---|---|---|---|---|---|---|
| | | | ΔG° (pH 7.0) | ΔG°′ (pH 5.5) | ΔG° (pH 7.0) | ΔG°′ (pH 5.5) | ΔG° (pH 7.0) | ΔG°′ (pH 5.5) | ΔG° (pH 7.0) | ΔG°′ (pH 5.5) |
| | | **Methane production** | | | | | | | | |
| 1 | Hydrogenotrophic methanogenesis | $4H_2 + CO_2 \rightarrow CH_4 + 2H_2O$ | -130.73 | -130.35 | -128.69 | -128.71 | -123.82 | -123.75 | -116.88 | -116.78 |
| 2 | Acetoclastic methanogenesis | $CH_3COO^- + H^+ \rightarrow CH_4 + CO_2$ | -35.69 | -138.4 | -36.61 | -140.1 | -38.17 | -144.24 | -40.37 | -150.06 |
| | | **Acetate production** | | | | | | | | |
| 3 | Ethanol oxidation | $CH_3CH_2OH + H_2O \rightarrow CH_3CHOO^- + H^+ + 2H_2$ | 9.89 | 112.14 | 8.33 | 111.84 | 4.62 | 110.64 | -0.65 | 108.95 |
| 4 | Homoacetogenesis in *C. thermoaceticum* | $4H_2 + 2CO_2 \rightarrow CH_3COO^- + H^+ + 2H_2O$ | -94.77 | 8.05 | -92.08 | 11.39 | -85.65 | 20.49 | -76.51 | 33.28 |
| 5 | Lactate oxidation | $CH_3CH(OH)COO^- + H_2O \rightarrow CH_3COO^- + 2H_2 + CO_2$ | -9.47 | 52.7 | -12.09 | 51.7 | -18.35 | 47.91 | -27.24 | 42.55 |
| | | **Propionate production** | | | | | | | | |
| 6 | Lactate reduction to propionate: as found in *Selenomonas ruminantium* | $CH_3CH(OH)COO^- + H_2O \rightarrow CH_3COO^- + CO_2 + 2H_2$ $CH_3CH(OH)COO^- + H_2 \rightarrow CH_3CH_2COO^- + H_2O$ ×2 | -171.87 | 15.46 | -175.47 | 17.84 | -184.13 | 14.77 | -196.38 | 13.19 |
| 7 | Lactate reduction to propionate: as determined for *C. propionicum* | $CH_3CH(OH)COO^- + H_2 \rightarrow CH_3CH_2COO^- + H_2O$ | -81.2 | -18.62 | -81.69 | -16.93 | -82.89 | -16.57 | -84.57 | -14.68 |
| 8 | Propionate formation in *Pelobacter propionicus* | $3CH_3CH_2OH + 2CO_2 \rightarrow 2CH_3CH_2COO^- + CH_3COO^- + H_2O + 3H^+$ | -113.84 | 193.54 | -114.24 | 197.27 | -115.14 | 202.88 | -115.86 | 181.15 |
| | | **Chain elongation to even-chain products with ethanol** | | | | | | | | |
| 9 | Ethanol to *n*-butyrate | $CH_3CH_2OH + CH_3COO^- \rightarrow CH_3(CH_2)_2COO^- + H_2O$ | -38.6 | -38.8 | -38.59 | -38.6 | -38.6 | -38.59 | -38.58 | -38.58 |
| 10 | Ethanol to *n*-caproate | $CH_3CH_2OH + CH_3(CH_2)_2COO^- \rightarrow CH_3(CH_2)_4COO^- + H_2O$ | -38.8 | -38.2 | -38.8 | -38.8 | -38.81 | -38.8 | -38.8 | -38.8 |
| 11 | Ethanol to *n*-caprylate | $CH_3CH_2OH + CH_3(CH_2)_4COO^- \rightarrow CH_3(CH_2)_6COO^- + H_2O$ | -43 | -43.24 | -43.37 | -43.32 | -43.51 | -43.8 | -43.4 | -43.03 |
| | | **Chain elongation odd-chain products with ethanol** | | | | | | | | |
| 12 | Ethanol to *n*-valerate | $CH_3CH_2OH + CH_3CH_2COO^- \rightarrow CH_3(CH_2)_3COO^- + H_2O$ | -38.60 | -38.43 | -38.60 | -39.59 | -38.60 | -38.59 | -38.58 | -38.58 |
| 13 | Ethanol to *n*-heptanoate | $CH_3CH_2OH + CH_3(CH_2)_2COO^- \rightarrow CH_3(CH_2)_5COO^- + H_2O$ | -42.05 | -41.85 | -42.15 | -42.11 | -42.25 | -42.24 | -42.42 | -42.23 |
| | | **Chain elongation to even-chain products with lactate** | | | | | | | | |
| 14 | Lactate to *n*-butyrate | $CH_3CH(OH)COO^- + CH_3COO^- + H^+ \rightarrow CH_3(CH_2)_2COO^- + H_2O + CO_2$ | -57.96 | -98.24 | -59.01 | -98.74 | -61.57 | -101.32 | -65.17 | -104.98 |
| 15 | Lactate to *n*-caproate | $CH_3CH(OH)COO^- + CH_3(CH_2)_2COO^- + H^+ \rightarrow CH_3(CH_2)_4COO^- + H_2O + CO_2$ | -58.16 | -97.64 | -59.22 | -98.94 | -61.78 | -101.53 | -65.39 | -105.2 |
| 16 | Lactate to *n*-caprylate | $CH_3CH(OH)COO^- + CH_3(CH_2)_4COO^- + H^+ \rightarrow CH_3(CH_2)_6COO^- + H_2O + CO_2$ | -62.60 | -102.3 | -63.79 | -105.3 | -66.47 | -106.23 | -70.34 | -110.09 |
| | | **Chain elongation to odd-chain products with lactate** | | | | | | | | |
| 17 | Lactate to *n*-valerate | $CH_3CH(OH)COO^- + CH_3CH_2COO^- + H^+ \rightarrow CH_3(CH_2)_3COO^- + H_2O + CO_2$ | -57.96 | -97.87 | -59.02 | -99.73 | -61.57 | -101.32 | -65.17 | -104.98 |
| 18 | Lactate to *n*-heptanoate | $CH_3CH(OH)COO^- + CH_3(CH_2)_2COO^- + H^+ \rightarrow CH_3(CH_2)_5COO^- + H_2O + CO_2$ | -61.41 | -101.29 | -62.57 | -102.25 | -65.22 | -104.97 | -68.01 | -108.83 |
| | | **ß-oxidation of carboxylic acids** | | | | | | | | |

| | | | | | | | | | | |
|---|---|---|---|---|---|---|---|---|---|---|
| 19 | *n*-caprylate to *n*-caproate | $CH_3(CH_2)_6COO^- + 2H_2O \rightarrow CH_3(CH_2)_4COO^- + CH_3COO^- + 2H_2 + H^+$ | 53.13 | 155.58 | 51.55 | 155.04 | 47.56 | 154.18 | 43.00 | 152.78 |
| 20 | *n*-caproate to *n*-butyrate | $CH_3(CH_2)_4COO^- + 2H_2O \rightarrow CH_3(CH_2)_2COO^- + CH_3COO^- + 2H_2 + H^+$ | 48.69 | 151.14 | 47.03 | 150.52 | 42.87 | 149.49 | 38.05 | 147.83 |
| 21 | *n*-butyrate to acetate | $CH_3(CH_2)_2COO^- + 2H_2O \rightarrow 2CH_3COO^- + 2H_2 + H^+$ | 48.49 | 150.94 | 46.83 | 150.32 | 42.66 | 149.28 | 37.83 | 147.61 |

[a] Adapted from (Cavalcante, Leitão *et al*. 2017). $\Delta G^{\circ}$: Gibbs free energy at different temperatures with 1M of substrates and products, a water activity of 1, gas partial pressure of 105 KPa, and pH=7. $\Delta G^{\circ\prime}$: Gibbs free energy at different temperatures with ion concentrations of 0 M, substrate and product concentrations of 1 M, a water activity of 1, gas partial pressures of 105 KPa, and pH=5.5.

**Table S9.** The average concentration of carboxylates in the bioreactor with a functional or dysfunctional extraction system in R1 and R2.

| ($mmol\ L^{-1}$) | Acetate | Propionate | *n*-Butyrate | *n*-Valerate | *n*-Caproate | *n*-Heptanoate | *n*-Caprylate | *n*-Nonanoate |
|---|---|---|---|---|---|---|---|---|
| **With a functional extraction system** | | | | | | | | |
| R1 | 8.81±0.12 | 0.90±0.22 | 10.6±0.51 | 1.50±0.15 | 12.5±0.44 | 0.07±0.01 | 1.95±0.15 | 0.00 |
| R2 | 3.94±0.06 | 1.01±0.04 | 6.60±0.20 | 1.40±0.01 | 8.30±0.33 | 0.20±0.02 | 1.34±0.16 | 0.00 |
| **With a dysfunctional extraction system** | | | | | | | | |
| R1 (Period V) | 3.11±0.33 | 4.14±0.17 | 20.9±0.48 | 0.12±0.02 | 20.0±0.27 | 0.13±0.03 | 4.39±0.11 | 0.00 |
| R2 (Period **b**) | 0 | 0.78±0.14 | 21.2±0.98 | 0.80±0.13 | 20.4±0.54 | 0.21±0.08 | 4.10±0.05 | 0.00 |
